# Membrane progesterone receptors spatially organize and partner with ABHD2 to mediate metabolic signaling

**DOI:** 10.64898/2026.11.19.766123

**Authors:** Shereena Sherif, Lama Assaf, Raghad El Ghali, Mariam Afifi, Mustafa Shammout, Khaled Machaca, Nancy Nader

## Abstract

Membrane progesterone receptors (mPRs) mediate rapid hormone responses, yet how they engage effectors remains poorly understood. Here, we reveal a spatially regulated partnership between mPRs and the lipid hydrolase ABHD2 that provides a mechanistic basis for the early steps of non- genomic progesterone signaling in mammalian cells. Agonist stimulation promotes mPRα- dependent accumulation of predominantly intracellular ABHD2 at the plasma membrane and enhances its association with mPRs. Although human ABHD2 binds progesterone, its phospholipase A₂ activity requires partnership with mPRs and an intact catalytic site. ABHD2 deletion disrupts agonist-induced metabolic signaling in hepatic and muscle cells and affects glucose homeostasis in obese mice. Together, these findings uncover a conserved receptor- hydrolase signaling mechanism in which spatially regulated association and receptor-dependent enzymatic activity emerge as defining features of non-genomic steroid signaling.

**Significance Statement:** We reveal a spatially regulated partnership between human mPRs and ABHD2 that enables lipase activation during early steps of rapid progesterone signaling and establish its functional significance in mammalian metabolic regulation.

## Introduction

Progesterone (P4) regulates diverse physiological processes through both genomic and non- genomic mechanisms^1–5^. Its classical actions are mediated by nuclear progesterone receptors (nPRs), ligand-activated transcription factors that regulate gene expression^4^. P4 also elicits rapid, non-genomic responses through several membrane-associated receptors, among which are membrane progesterone receptors (mPRs), members of the progestin and adipoQ receptor (PAQR) family^6^. Five mPR subtypes have been identified: mPRα (PAQR7), mPRβ (PAQR8), mPRγ (PAQR5), mPRδ (PAQR6) and mPRε (PAQR9). Expressed across reproductive and non- reproductive tissues, mPRs have been implicated in oocyte maturation, neuroprotection, inflammatory responses, tumor biology and glucose homeostasis^6–10^. Despite this functional diversity, the early molecular events coupling mPRs activation to downstream cellular responses remain poorly understood.

Our work in *Xenopus laevis* oocytes identified the lipid hydrolase α/β hydrolase domain-containing protein 2 (ABHD2) as an essential partner in mPRβ signaling^11^. ABHD2 was previously identified as a P4-binding protein in human sperm, where its monoacylglycerol lipase activity mediates rapid progesterone signaling by relieving lipid-dependent inhibition of the CatSper channel^12^. In oocytes, however, P4 enhances ABHD2 association with mPRβ and stimulates phospholipase A₂ (PLA₂) activity, which requires both proteins and is necessary for meiotic maturation^11^. Our earlier work also demonstrated that mPRβ endocytosis enables recruitment of the adaptor protein APPL1 and the kinase AKT2 to signaling endosomes^11, 13^. These findings identify receptor-enzyme partnership and regulated trafficking as linked components of progesterone-induced oocyte maturation.

Whether the mPR-ABHD2 partnership represents a conserved mechanism of progestin signaling beyond *Xenopus* oocytes remains unknown. The subcellular sites and mechanisms regulating their association are also undefined, as is the requirement for ABHD2 in mPR-dependent cellular responses in mammalian cells. Resolving these questions would establish the broader relevance of this partnership and reveal how membrane steroid receptors engage enzymatic effectors during rapid, non-genomic signaling.

The metabolic actions of mPRs provide a physiologically relevant setting to assess the functional significance of this partnership. We recently showed that Org OD 02-0 (OD), a progestin analogue reported to selectively activate mPRs over nPRs^14^, stimulates AMPK signaling and glucose uptake in mammalian cells and improves glucose homeostasis in obese mice^8^. Whether these responses require ABHD2, however, remains unknown.

Here, we investigate the role of ABHD2 in mPR-mediated signaling and metabolic regulation in mammalian cells and mice. We show that agonist stimulation promotes mPRα-dependent recruitment of ABHD2 to the plasma membrane and its association with mPRs, while human mPRα/β enable ABHD2-dependent PLA₂ activity. ABHD2 is required for mPRs agonist-induced metabolic responses in cells and contributes to glucose lowering in obese mice. Together, these findings reveal a conserved, spatially regulated receptor-hydrolase partnership linking rapid steroid signaling to metabolic control.

## Results

### Agonist stimulation promotes mPRα-dependent accumulation of ABHD2 at the plasma membrane

To investigate the subcellular localization of ABHD2, we first generated ABHD2-knockout HepG2 cells using CRISPR/Cas9, providing a defined background for imaging tagged ABHD2 in the absence of the endogenous protein. Sequencing of the targeted region identified mutations in two isolated clones, T2-32 and T2-44, with the predicted amino acid sequence of clone T2-44 containing premature stop codons (Supplementary Fig. 1A, B). Clone T2-44 was used for all experiments involving ABHD2-knockout HepG2 cells and is hereafter referred to as ABHD2-KO. We then performed high-resolution confocal imaging in ABHD2-KO cells re-expressing ABHD2- His together with the plasma membrane marker Orai1 (Fig. 1A). Although ABHD2 has been described as a plasma membrane progesterone receptor¹², it was predominantly intracellular under basal conditions in HepG2 cells, with limited plasma membrane localization. OD stimulation for 30 min promoted ABHD2 accumulation at the cell periphery, where its fluorescence overlapped with Orai1 (Fig. 1A). Fluorescence intensity profiles supported this redistribution, revealing a significant reduction in the distance between ABHD2 and Orai1 fluorescence peaks (Fig. 1B, C).

**Figure 1.**
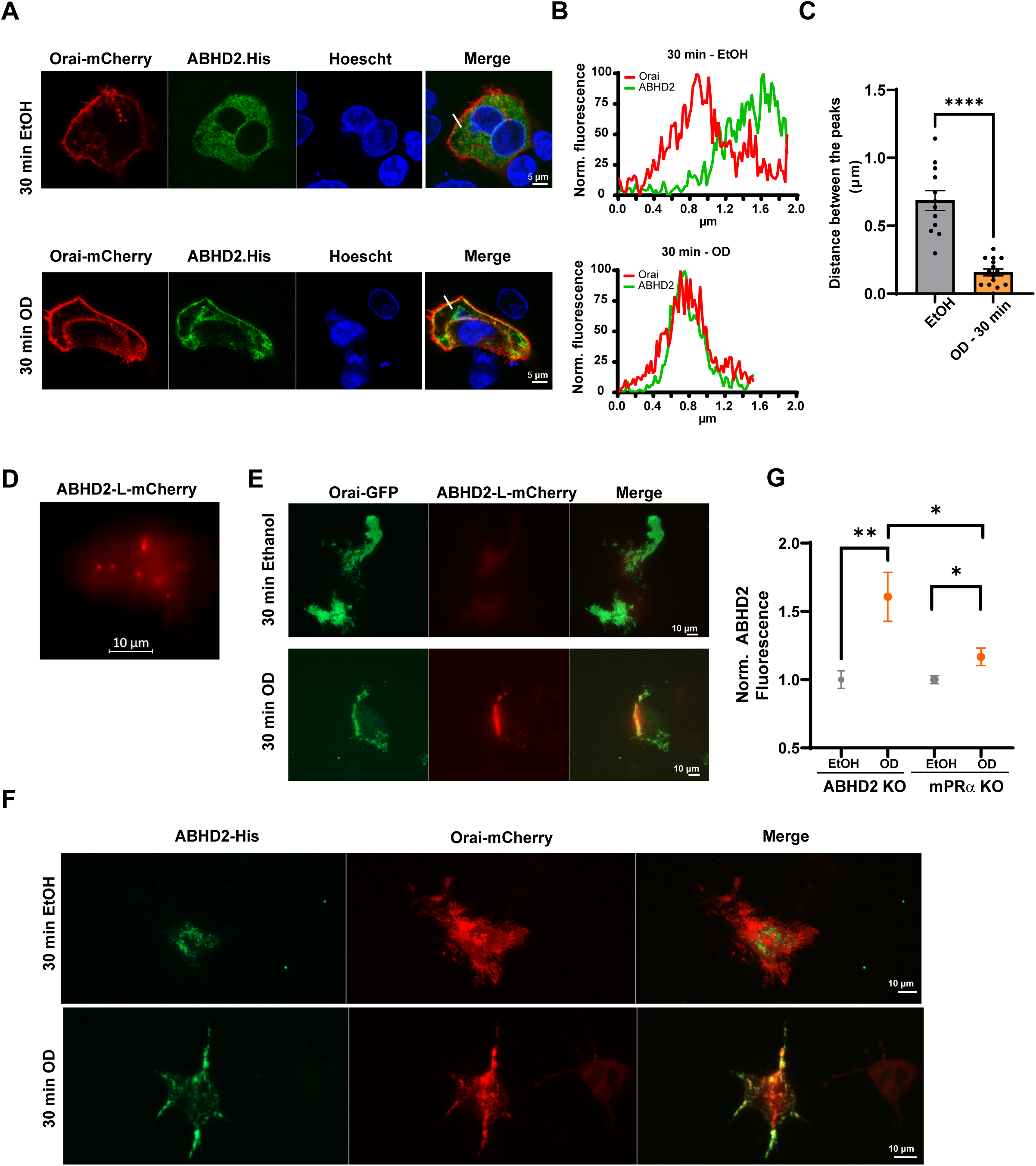
Agonist stimulation promotes mPRα-dependent accumulation of ABHD2 near the plasma membrane. **(A)** Representative confocal images of ABHD2-KO HepG2 cells expressing ABHD2-His and Orai1-mCherry after treatment with EtOH or OD for 30 min. ABHD2-His was detected by anti- His immunostaining (green), Orai1-mCherry marks the plasma membrane (red), and nuclei were stained with Hoechst (blue). White lines indicate the regions used for fluorescence line-scan analysis. Scale bar, 5 μm. **(B)** Representative normalized fluorescence intensity profiles of Orai1-mCherry and ABHD2-His along the lines shown in (A). Each fluorescence profile was normalized to its own maximum intensity. **(C)** Quantification of the absolute distance between the Orai1 and ABHD2 fluorescence peaks. Each point represents a line scan from a single cell. Data are presented as mean ± SEM from 12 to 15 cells from 3 independent experiments. Ethanol and OD values were compared using unpaired *t* test. **(D)** Representative frame from a live-cell total internal reflection fluorescence (TIRF) recording showing dynamic ABHD2-L-mCherry-positive structures near the plasma membrane under basal conditions. See Supplementary Video 1. Scale bar, 10 μm. **(E)** Representative TIRF images of ABHD2-KO HepG2 cells co-expressing ABHD2-L-mCherry (red) and Orai1-GFP (green), following EtOH or OD treatment for 30 min. Scale bar, 10 μm. **(F)** Representative TIRF images of ABHD2-KO HepG2 cells expressing ABHD2-His (green) and Orai1-mCherry (red), following EtOH or OD treatment for 30 min. Scale bar, 10 μm. **(G)** Quantification of ABHD2 fluorescence in the TIRF imaging plane in ABHD2-KO and mPRα- KO cells re-expressing ABHD2-His along with Orai-mCherry. Representative mPRα-KO images are shown in Fig. S2G. Fluorescence was normalized to the corresponding EtOH condition. Data are presented as mean ± SEM from 39 to 49 cells from 3 independent experiments. Ethanol and OD values were compared using unpaired *t* test. For all experiments, the OD concentration used is 10^-6^M, and Ethanol at 0.1%. Statistical significance: *p < 0.05; **p < 0.01; ****p < 0.0001.

To examine ABHD2 dynamics near the plasma membrane, we performed live-cell total internal reflection fluorescence (TIRF) microscopy using ABHD2 tagged to mCherry through a flexible linker (ABHD2-L-mCherry; Supplementary Fig. 1C). The linker was introduced because direct fusion of ABHD2 to a fluorescent protein previously impaired its function^11^. Under basal conditions, ABHD2-positive structures dynamically entered and exited the evanescent field, consistent with cycling movement toward and away from the plasma membrane (Fig. 1D and Supplementary Video 1A, B). ABHD2-positive puncta and elongated, track-like structures were also observed near the cell surface (Supplementary Fig. 1D), with dynamic movements evident in live-cell imaging (Supplementary Video 2). Following OD stimulation, live-cell imaging showed increased membrane-proximal ABHD2 fluorescence and overlap with Orai1-GFP (Fig. 1E). To quantify this response across cells, we performed fixed-cell TIRF imaging after vehicle or 30-min OD treatment by re-expressing ABHD2-His. This analysis confirmed OD-induced ABHD2 accumulation near the plasma membrane (Fig. 1F, G), consistent with the behavior observed using ABHD2-L-mCherry.

We next investigated whether OD-induced ABHD2 accumulation at the plasma membrane depends on mPRα, given its reported expression in human liver and functional involvement in progestin signaling in HepG2 cells^15^. To this end, we generated mPRα-knockout HepG2 cells using CRISPR/Cas9. Targeted mutations in two independent clones (T2-15 and T2-16) were confirmed by sequencing, and loss of mPRα protein was verified by immunoblotting (Supplementary Fig. 2A-C). Clone T2-15 was used for all experiments involving mPRα-knockout HepG2 cells and is hereafter referred to as mPRα-KO. Before examining its role in ABHD2 redistribution, we first verified that mPRα mediates the metabolic responses previously observed following OD stimulation in HepG2 cells^8^. mPRα deletion abolished OD-induced AMPK phosphorylation (Supplementary Fig. 2D). In this clone, OD reduced rather than increased intracellular glucose accumulation, as measured using a genetically encoded FRET sensor (Supplementary Fig. 2E, F). These findings established mPRα as a functionally relevant receptor for investigating the link between ABHD2 redistribution and metabolic signaling. TIRF imaging then revealed markedly reduced OD-induced ABHD2 accumulation at the plasma membrane in mPRα-knockout cells (Fig. 1G and Supplementary Fig. 2G).

Together, these findings show that ABHD2 dynamically accesses the plasma membrane and identify mPRα as a regulator of its spatial distribution, supporting agonist-induced accumulation at the cell surface.

## Human ABHD2 binds progesterone

The predominantly intracellular localization of ABHD2 prompted us to examine its role in hormone recognition. Membrane-impermeant P4-BSA can initiate signaling at the cell surface^5^, where ABHD2 is only transiently present under basal conditions. We therefore asked whether human ABHD2 retains the ability to bind P4, as previously reported^12^.

To address this question, we performed P4-biotin pull-down assays using recombinant human His- tagged ABHD2 and mPRα expressed in the pALiCE system (Supplementary Fig. 3A). Proteins were incubated with or without P4-biotin, captured on streptavidin beads, eluted and analyzed by Western blotting. Human mPRα served as a positive control and was detected in the pull-down fraction following incubation with P4-biotin. After subsequent incubation with excess free P4 or OD, mPRα was no longer detectable in the bead-bound fraction. ABHD2 showed a similar pattern, consistent with displacement of both proteins from P4-biotin by free P4 or OD (Supplementary Fig. 3B). These findings support progesterone binding by human ABHD2 and are consistent with OD binding.

## Agonist stimulation promotes ABHD2 association with mPRs

The mPRα-dependent accumulation of ABHD2 near the plasma membrane prompted us to investigate whether agonist stimulation also promotes its association with mPRs. We first examined the distribution of mPRα-His and ABHD2-L-mCherry in HepG2 cells. Under basal conditions, ABHD2 was predominantly intracellular, with limited overlap with peripheral mPRα staining. OD stimulation for 30 min increased their colocalization at the cell periphery (Fig. 2A), supported by increased Pearson’s correlation coefficients within peripheral regions of interest (Fig. 2B).

**Figure 2.**
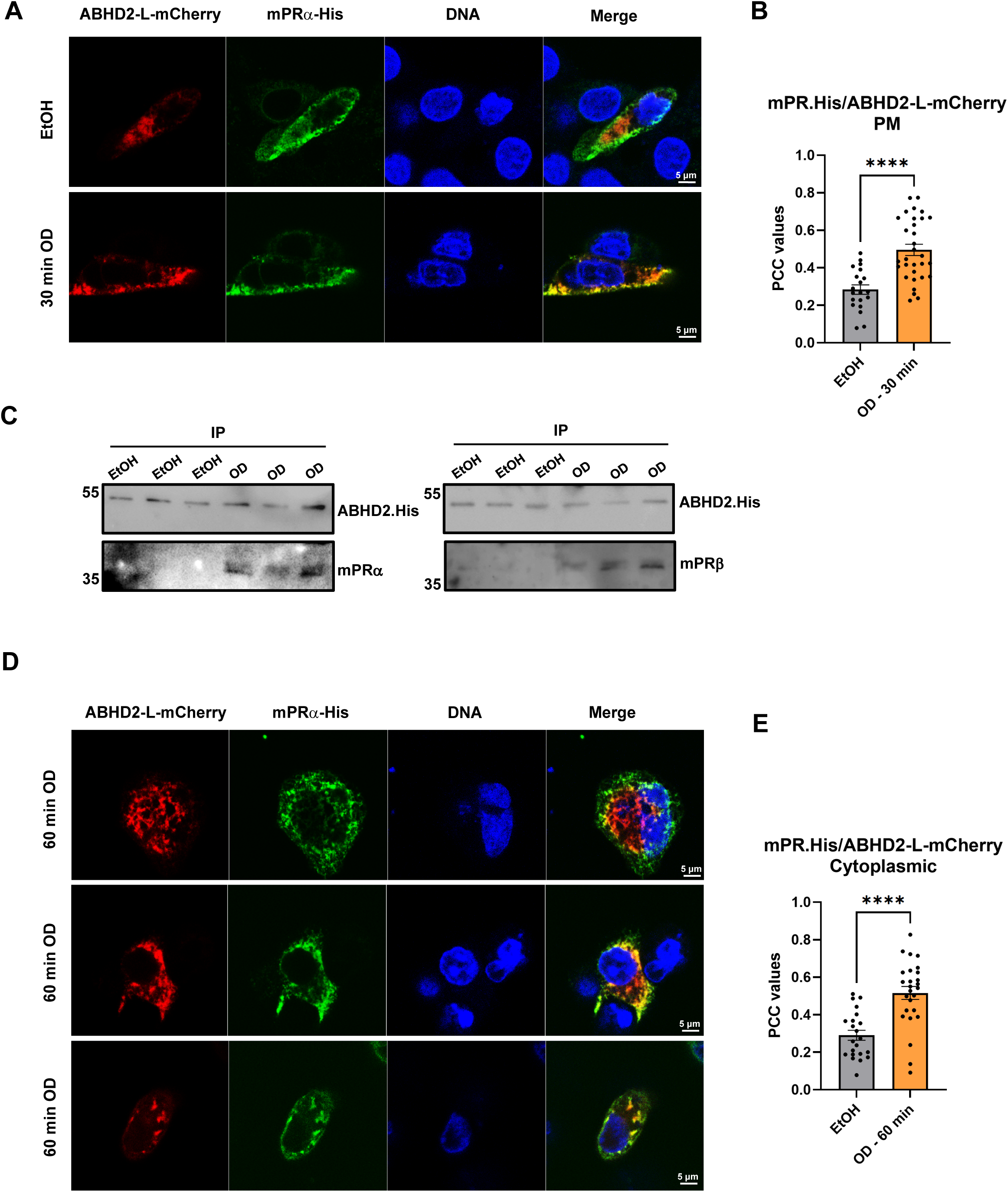
Agonist stimulation promotes ABHD2 association with mPRs **(A)** Representative confocal images of HepG2 cells co-expressing mPRα-His and ABHD2-L- mCherry following treatment with EtOH or OD (10⁻⁶ M) for 30 min. mPRα-His was detected by anti-His immunostaining, ABHD2-L-mCherry was visualized through mCherry fluorescence, and nuclei were stained with Hoechst (blue). Scale bar, 5 μm. **(B)** Quantification of mPRα-ABHD2 colocalization using Pearson’s correlation coefficient (PCC) within mPRα-defined regions of interest at the cell periphery. Data are presented as mean ± SEM; n = 20 cells (EtOH) and 31 cells (OD) from three independent experiments. EtOH- and OD-treated groups were compared using an unpaired *t* test. **(C)** His-tag pull-downs from ABHD2-KO HepG2 cells re-expressing ABHD2-His following treatment with EtOH or OD (10⁻⁶ M) for 30 min. Eluted fractions were immunoblotted for ABHD2-His and endogenous mPRα and mPRβ. The three EtOH and three OD lanes represent samples from three independent experiments. **(D)** Representative confocal images of HepG2 cells co-expressing mPRα-His and ABHD2-L- mCherry following OD treatment (10⁻⁶ M) for 60 min. Proteins and nuclei were visualized as described in (A). Scale bar, 5 μm. **(E)** Quantification of mPRα-ABHD2 colocalization using PCC within cytoplasmic regions of interest. Data are presented as mean ± SEM; n = 22 cells (EtOH) and 26 cells (OD) from three independent experiments. Multiple comparisons were done using an unpaired *t* test.

To determine whether this increased colocalization was accompanied by biochemical association, we performed pull-down assays in ABHD2-KO HepG2 cells expressing ABHD2-His following 30 min of vehicle or OD treatment. Endogenous mPRα was recovered with ABHD2-His following OD stimulation, with little or no recovery under vehicle conditions (Fig. 2C). Because mPRβ is also expressed in HepG2 cells (Supplementary Fig. 3C), we examined whether it similarly associates with ABHD2. Endogenous mPRβ likewise showed OD-enhanced recovery with ABHD2-His (Fig. 2C). These findings extend our previous observations in *Xenopus* oocytes^11^, demonstrating agonist-enhanced association of human ABHD2 with both mPRα and mPRβ in mammalian cells.

Given our previous finding that mPRβ undergoes endocytosis after approximately 1 h of P4 stimulation in *Xenopus* oocytes^11, 13^, we examined mPRα and ABHD2 localization after 60 min of OD treatment. The two proteins showed overlapping intracellular staining (Fig. 2D), accompanied by increased PCC within cytoplasmic regions of interest (Fig. 2E).

Together, these findings support agonist-enhanced ABHD2 association with mPRs and reveal a coordinated pattern of ABHD2-mPRα colocalization at the cell periphery at 30 min and within intracellular regions at 60 min.

## Human ABHD2 functions as an mPR-dependent phospholipase A₂

Having established agonist-enhanced association of ABHD2 with mPRs, we next examined the enzymatic consequences of this partnership. ABHD2 has been reported to exhibit monoacylglycerol lipase activity in human sperm^12^, whereas our previous work showed that ABHD2 functions as a phospholipase A₂ (PLA₂) in *Xenopus* oocytes^11^. To determine whether this enzymatic function is conserved in mammalian cells, we measured PLA₂ activity in HepG2 cell lysates following vehicle or OD treatment. Control cells exhibited basal PLA₂ activity that increased significantly after OD stimulation (Fig. 3A, D). This increase was lost in ABHD2- and mPRα-knockout cells (Fig. 3B-D), indicating that both proteins are required for the OD-induced PLA₂ response.

**Figure 3.**
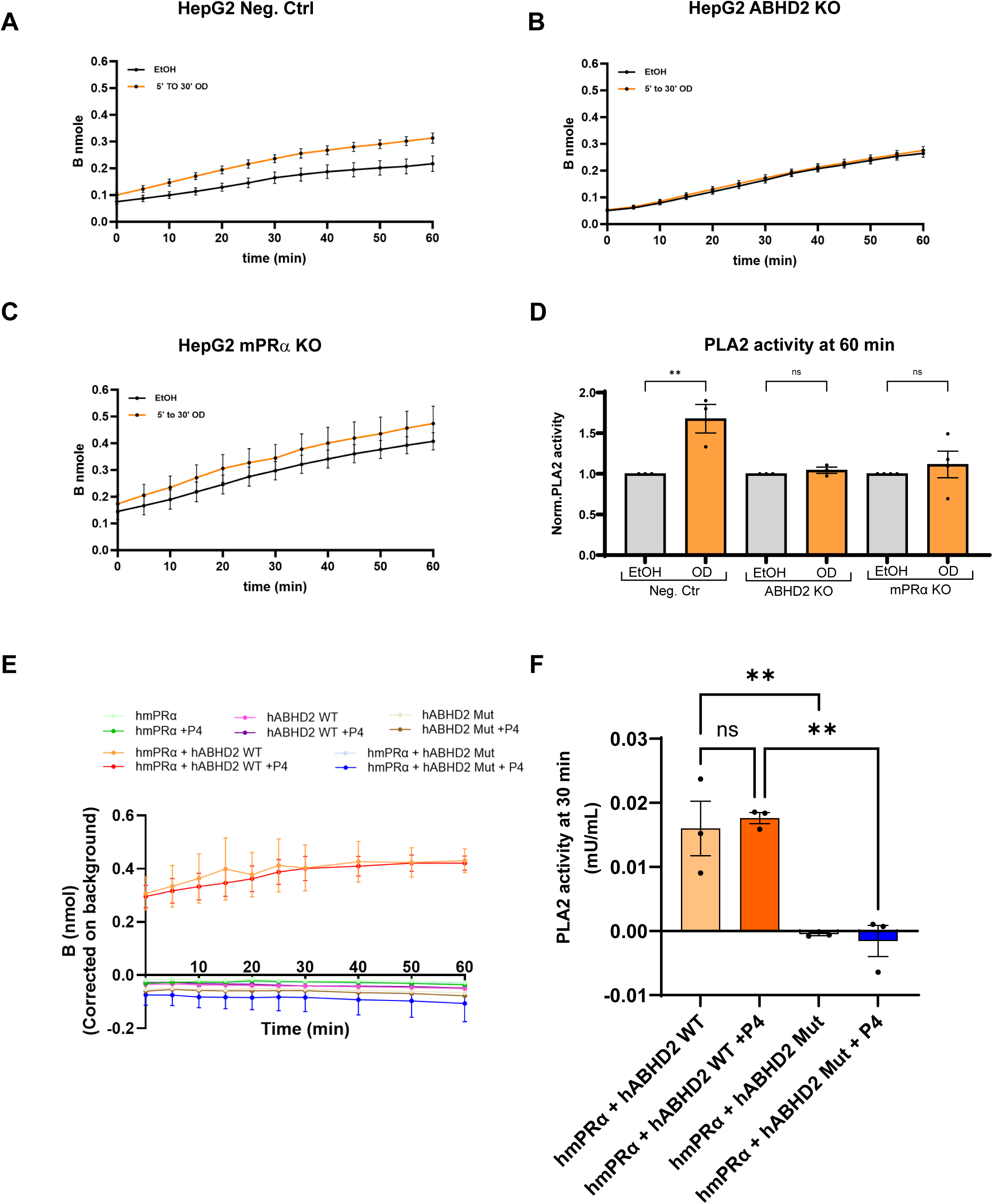
Human ABHD2-PLA2 activity is dependent on mPRαχσβαρλινε **(A-C)** Time courses of PLA₂ substrate hydrolysis in lysates from Negative control (Neg. Ctrl.) (A), ABHD2-KO (B), and mPRα-KO (C) HepG2 cells treated with EtOH (0.1%) or OD (10^-7^M). Cells were exposed to OD for 5 to 30 min before lysis, and fluorescence was monitored during the 60- min assay. The OD treatment time point yielding maximal activity in each independent experiment was plotted. Data are presented as mean ± SEM; n = 3 to 4 independent experiments per cell type. **(D)** Quantification of PLA₂ activity from the 60-min assay readout, normalized to the corresponding EtOH condition. Data are presented as mean ± SEM; n = 3 to 4 independent experiments per cell type. Statistical analysis was performed using ordinary one-way ANOVA followed by Tukey’s multiple-comparisons test. EtOH-versus-OD comparisons within each cell type are shown. **(E)** Time courses of background-corrected PLA₂ substrate hydrolysis in ALiCE microsomal preparations expressing human mPRα, WT ABHD2, or catalytic-site mutant ABHD2 (ABHD2 mut), individually or together as indicated, with or without P4 (10⁻⁵M). EtOH served as vehicle. Fluorescence was monitored for 60 min. Data are represented as mean ± SEM; n=3. **(F)** PLA₂ activity at 30 min in preparations co-expressing mPRα with WT or catalytic-site mutant ABHD2, in the presence or absence of P4. For D and E, Data are represented as mean ± SEM; n=3, one-way ANOVA was used for comparison. Statistical significance: **p < 0.01; ns, not significant.

To test whether this response requires ABHD2 catalytic activity rather than solely reflecting downstream lipase activation, we measured PLA₂ activity in a reconstituted system. Human ABHD2 and mPRα or mPRβ were expressed individually or together using the pALiCE cell-free expression system, with signal sequences directing the proteins to microsomal membranes. Expression of the receptors and wild-type or catalytic-site mutant ABHD2 was confirmed by immunoblotting (Supplementary Fig. 4A). Microsomal preparations containing ABHD2 alone or either receptor alone showed no detectable PLA₂ activity above background. In contrast, co- expression of wild-type ABHD2 with mPRα produced robust activity, which was lost when the receptor was co-expressed with the ABHD2 catalytic-site mutant (Fig. 3E, F). Co-expression with mPRβ similarly supported activity that was markedly reduced by mutation of the ABHD2 catalytic site (Supplementary Fig. 4B, C). Thus, both mPRα and mPRβ supported PLA₂ activity that depended on an intact ABHD2 catalytic site.

PLA₂ activity in microsomes co-expressing ABHD2 with either receptor was detectable without exogenous P4 and was not further enhanced by its addition (Fig. 3F and Supplementary Fig. 4C), consistent with our previous findings in *Xenopus laevis*^11^. This apparent ligand-independent activity may reflect elevated receptor abundance and confinement of the two proteins within the same membrane compartment, potentially bypassing the agonist-dependent spatial convergence observed in cells.

Together, these results support a conserved requirement for mPRs in ABHD2-associated PLA₂ activity.

## Endocytic trafficking contributes to mPR-mediated metabolic signaling

Our previous studies established a role for endocytosis in mPRβ signaling in *Xenopus* oocytes^11, 13, 16^. Live-cell TIRF imaging of mPRα-knockout HepG2 cells re-expressing mPRα-GFP revealed receptor localization at the plasma membrane, overlapping with Orai1-mCherry, together with dynamic receptor-positive structures near the cell surface (Supplementary Fig. 5A and Supplementary Video 3). These observations, together with the intracellular colocalization of ABHD2 and mPRα after 60 min of OD stimulation, prompted us to test whether endocytic trafficking contributes to OD-mediated metabolic signaling in mammalian cells.

We treated HepG2 cells with the endocytosis inhibitor Pitstop 2^17^, and verified its inhibitory effect by reduced transferrin uptake (Supplementary Fig. 5B). We then assessed AMPK phosphorylation, a previously established response to OD in HepG2 cells^8^. Following Pitstop 2 pretreatment, OD no longer significantly increases AMPK phosphorylation (Supplementary Fig. 5C, D). These observations are consistent with a contribution of endocytic trafficking to mPRs agonist-induced metabolic signaling.

## ABHD2 is required for mPR-induced metabolic responses in liver and muscle cells

Having established ABHD2 association with mPRs, receptor-dependent PLA₂ activity and a contribution of endocytic trafficking to metabolic signaling, we next asked whether ABHD2 is required for the metabolic effects of mPRs activation. We assessed responses characterized in our recent study^8^ : OD-induced AMPK activation in hepatic and muscle cells, and increased glycolysis and glucose uptake in HepG2 cells.

In HepG2 cells, OD markedly increased AMPK phosphorylation in controls, whereas no significant increase was detected in ABHD2-KO cells (Fig. 4A, B). Under vehicle conditions, ABHD2-KO cells showed higher mean basal AMPK phosphorylation, although this difference was variable and did not reach statistical significance (Supplementary Fig. 6A). OD also enhanced glycolysis in control cells but had no significant effect in ABHD2-KO cells (Fig. 4C, D). Re- expression of human ABHD2-L-mCherry restored this response, supporting the specificity of the knockout phenotype, whereas the ABHD2 catalytic-site mutant failed to rescue it (Fig. 4E, F). Together with the cellular and microsomal PLA₂ assays, these results support a role for ABHD2- dependent PLA₂ activity upstream of the glycolytic response to mPRs activation. Also, OD- induced glucose uptake was likewise lost in ABHD2-KO cells (Fig. 4G, H). Notably, basal glycolytic activity and glucose uptake were significantly higher in ABHD2-KO cells than in controls (Supplementary Fig. 6B, C), indicating that ABHD2 deletion has distinct effects on basal glucose metabolism and OD-induced metabolic signaling.

**Figure 4.**
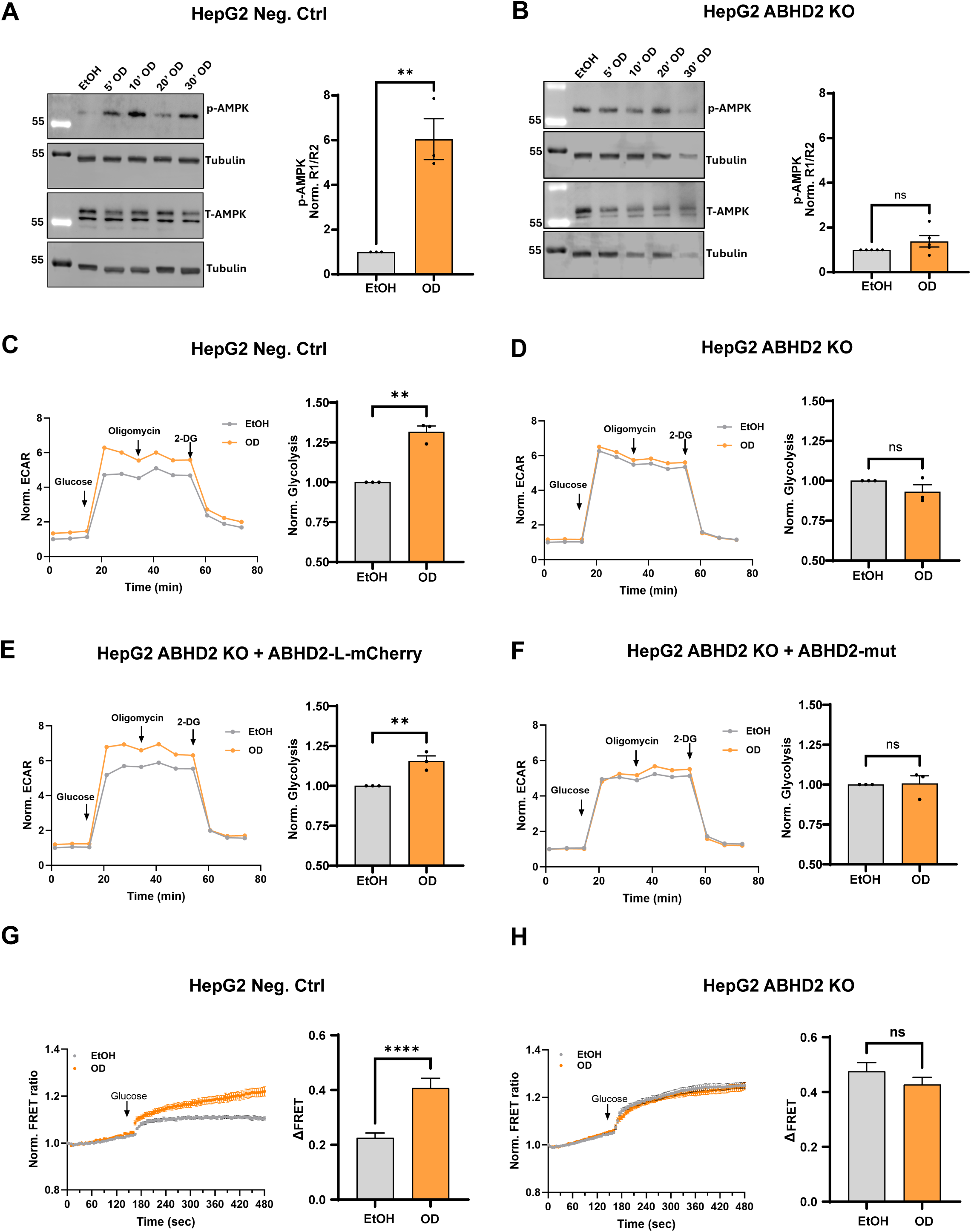
ABHD2 is required for OD-induced metabolic responses **(A, B)** Representative immunoblots (left) and quantification (right) of AMPK phosphorylation in negative control (Neg. Ctrl) (A) and ABHD2-KO (B) HepG2 cells treated with EtOH (0.1%) or OD (10⁻⁷ M) for 5, 10, 20, or 30 min. Tubulin served as the loading control. Densitometric quantification was performed using ImageJ. For each sample, phospho-AMPK/Tubulin (R1) and total-AMPK/Tubulin (R2) ratios were calculated, and phosphorylation was expressed as R1/R2 and normalized to EtOH (Norm. R1/R2). The OD treatment time point yielding maximal activity in each independent experiment was plotted. Data are presented as mean ± SEM; n= 3 to 5 independent experiments. Before- and after-treatment values were compared using unpaired *t* test. **(C, D, E, F)** Representative extracellular acidification rate (ECAR) traces (left) and quantification of glycolysis across all three independent experiments (right) in negative-control HepG2 cells (C), ABHD2-KO HepG2 cells (D), and ABHD2-KO HepG2 cells re-expressing ABHD2-L-mCherry (E) or the His-tagged ABHD2 catalytic-site mutant (F), treated with EtOH (0.1%) or OD (10⁻⁶ M). Glycolytic function was assessed using the Seahorse XF Glycolysis Stress Test. Arrows indicate sequential injections of glucose, oligomycin, and 2-deoxyglucose (2-DG). ECAR and glycolysis values were normalized to the corresponding vehicle condition (0.1% ethanol) within each experiment. Glycolysis data are presented as mean ± SEM across three independent experiments. Vehicle- and OD-treated groups were compared using an unpaired *t* test. **(G, H)** Time courses of normalized YFP/CFP FRET ratio (left) and quantification of raw ΔFRET (right) in negative control (Neg. Ctrl) (G) and ABHD2-KO (H) in HepG2 cells expressing the FLII¹²Pglu-700μδ6 glucose sensor. Cells were pretreated with EtOH (0.1%) or OD (10⁻^6^M) for 30 min before addition of 1 mM glucose at the indicated time. Ratios were normalized to the pre- glucose baseline. ΔFRET were calculated as the increase in the YFP/CFP ratio without normalization after glucose addition relative to the ratio immediately before addition. Data are presented as mean ± SEM; n = 32 to 42 cells per condition from 3 independent experiments. Values were compared using unpaired *t* test. Statistical significance: **p < 0.01; ****p < 0.0001; ns, not significant.

To determine whether the requirement for ABHD2 extends to muscle cells, we generated ABHD2- knockout C2C12 cells, with targeted mutations confirmed in two independent clones T2-8 and T2- 16 (Supplementary Fig. 6D, E). All subsequent experiments in ABHD2-knockout C2C12 cells used clone T2-16, designated ABHD2-KO. OD increased AMPK phosphorylation in control cells, but this response was lost following ABHD2 deletion (Supplementary Fig. 6F, G).

Together, these findings establish a requirement for ABHD2 in mPRs agonist-induced metabolic signaling in hepatic and muscle cells. They extend the functional relevance of the mPR-ABHD2 partnership beyond *Xenopus laevis* oocytes^11^, supporting conservation of the mPR-ABHD2 signaling module across vertebrates.

## ABHD2 contributes to the glucose-lowering response to OD in obese mice

Having established that ABHD2 is required for mPRs agonist-induced metabolic signaling in hepatic and muscle cells, we next asked whether it is also required for OD-induced glucose lowering in mice with diet-induced obesity and hyperglycemia^8^. We first characterized ABHD2- knockout (KO) mice and C57BL/6N controls under chow-fed conditions and following high-fat diet (HFD) feeding. Chow-fed ABHD2 KO mice exhibited modestly but significantly lower fasting glucose levels despite slightly higher body weight than controls (Supplementary Fig. 7A, D). Together with the elevated basal glycolytic activity and glucose uptake observed in ABHD2- deficient HepG2 cells, this finding suggests a role for ABHD2 in regulating basal glucose homeostasis. HFD feeding increased fasting glucose and body weight in both genotypes (Supplementary Fig. 7B, C, E, F).

We then treated HFD-fed ABHD2 KO mice with OD or vehicle for four weeks, monitoring body weight weekly (Supplementary Fig. 7J). Before treatment, fasting glucose was elevated relative to chow-fed controls and comparable between the HFD groups assigned to vehicle or OD (Fig. 5A). After treatment, fasting glucose remained elevated in both groups, with no significant difference between vehicle- and OD-treated ABHD2 KO mice (Fig. 5B). We therefore directly compared OD-associated changes in fasting glucose between C57BL/6N and ABHD2 KO mice. OD significantly reduced fasting glucose in C57BL/6N controls but not in ABHD2 KO mice (Supplementary Fig. 7G, H). The change in fasting glucose differed significantly between genotypes, demonstrating attenuation of the glucose-lowering response following ABHD2 deletion (Supplementary Fig. 7I). Individual KO mice showed variable responses, with fasting glucose decreasing in some animals and increasing in others (Supplementary Fig. 7H, I).

**Figure 5.**
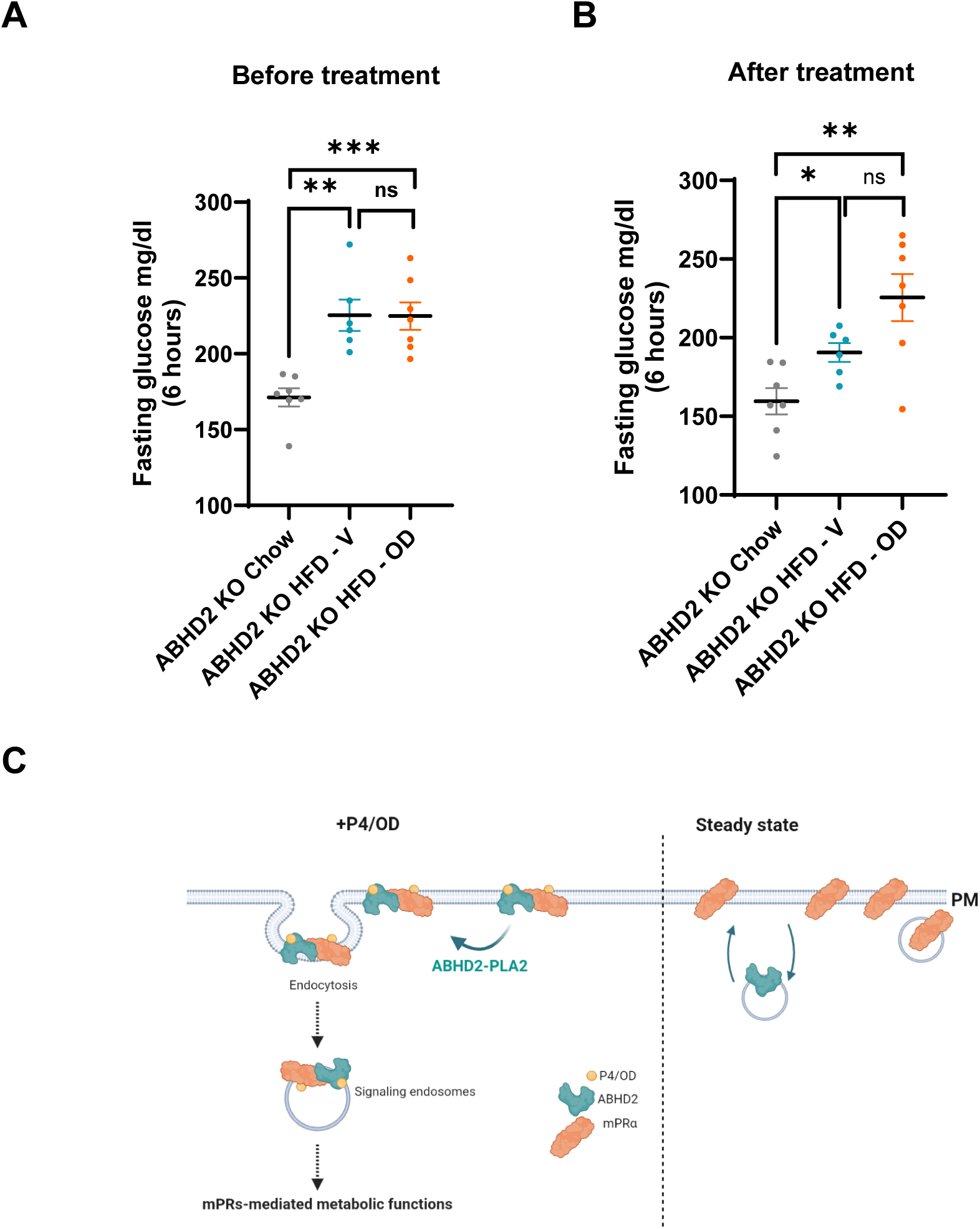
ABHD2 contributes to glucose regulation in mice and proposed model of spatially regulated mPR-ABHD2 signaling. **(A, B)** Blood glucose after a 6-h fast before (A) and after (B) treatment in chow-fed and HFD-fed ABHD2-KO mice. Each point represents one mouse. Data are presented as mean ± SEM; n=7 mice chow, n=6 mice HFD-V, n=7 mice HFD-OD. The statistical analysis was done using the Welch’s *t* test. Statistical significance: *p < 0.05; **p < 0.01; ***p < 0.001; ns, not significant. **(C)** Proposed model of spatially regulated mPR-ABHD2 signaling. Under basal conditions, ABHD2 resides predominantly in intracellular compartments and dynamically samples the membrane-proximal region. Agonist stimulation promotes mPRα-dependent accumulation of ABHD2 near the plasma membrane and enhances receptor-enzyme association. This partnership supports ABHD2-dependent PLA₂ activity, while endocytic trafficking is proposed to contribute to downstream metabolic signaling. Together, these findings support a model in which spatially regulated receptor-hydrolase assembly coordinates mPR-mediated metabolic responses. Created in BioRender. Nader, N. (2026) https://BioRender.com/jdl3ttj.

Together, these findings establish that ABHD2 contributes to OD-induced glucose lowering in obese mice, extending the functional relevance of the mPR-ABHD2 partnership to whole-body metabolic regulation.

## Discussion

Progesterone regulates diverse physiological processes through both nuclear receptor-dependent transcription and rapid membrane-initiated signaling^3, 6, 7, 9, 18–21^. Despite increasing recognition of these non-genomic actions, how membrane progesterone receptors couple steroid recognition to intracellular signaling remains not fully understood. Defining these events is complicated by the diversity of proteins implicated in rapid progesterone signaling ^6^, including mPRs and ABHD2, whose signaling functions have largely been investigated separately. Our previous work in *Xenopus* oocytes identified mPRβ and ABHD2 as obligate partners in progesterone-induced meiotic maturation^11^, raising the question of whether this partnership is restricted to oocyte maturation or operates more broadly in mammalian signaling.

Here, we establish a spatially regulated partnership between human mPRs and ABHD2 and demonstrate its relevance to metabolic regulation. Agonist stimulation promotes mPRα-dependent accumulation of predominantly intracellular ABHD2 at the plasma membrane and enhances its association with mPRα and mPRβ. Co-expression with either receptor supports PLA₂ activity requiring an intact ABHD2 catalytic site. ABHD2 deletion also disrupts agonist-induced metabolic responses in hepatic and muscle cells and attenuates OD-induced glucose lowering in obese mice (Fig. 5C). These findings extend the partnership beyond oocyte maturation and identify spatial regulation of ABHD2 as a feature of mammalian mPRs signaling. They also align with our parallel study in *Xenopus* oocytes^22^ regarding ABHD2 localization, progesterone binding and agonist- induced trafficking. Together, the two studies support conservation of this signaling module across vertebrates.

A central mechanistic finding is that mPRα regulates the spatial distribution of ABHD2. In HepG2 cells, ABHD2 was predominantly intracellular, with dynamic structures transiently accessing the membrane-proximal region. Its agonist-induced accumulation at the plasma membrane was substantially reduced following mPRα deletion, suggesting that the receptor promotes ABHD2 recruitment or retention at the cell surface. Both proteins bind progesterone, yet detectable PLA₂ activity in microsomal preparations required their co-expression and an intact ABHD2 catalytic site. Whereas OD enhanced PLA₂ activity in HepG2 cells, microsomal co-expression supported activity without exogenous progesterone. Confinement within a shared membrane compartment may facilitate productive receptor–hydrolase interactions, consistent with our parallel finding that increased receptor expression can overcome ligand dependence in *Xenopus*^22^. Together, these observations support a model in which agonist stimulation promotes spatial assembly of an enzymatically active mPR-ABHD2 complex. By regulating where ABHD2 encounters membrane- associated substrates, this unusual receptor-hydrolase partnership may provide spatial control over lipid hydrolysis and the initiation of downstream lipid signaling.

Our progesterone-binding findings support the original characterization of ABHD2 in human sperm^12^, although they differ from the absence of a detectable progesterone-induced thermal shift reported recently by Edwards et al.^23^. This discrepancy may reflect differences in protein preparation: ABHD2 remained associated with microsomal membranes in our preparations, whereas the other study used detergent-extracted, purified protein. The assays also measure different properties: ligand-associated protein recovery versus changes in protein stability. The molecular environment may likewise influence ABHD2 enzymatic activity. Although characterized as a monoacylglycerol lipase in sperm^12^, ABHD2 supports mPR-dependent PLA₂ activity in our mammalian preparations, extending our observations in *Xenopus* oocytes^11^. The ability of the related hydrolase ABHD6 to act on different lipid substrates^24^ raises the possibility that receptor association or membrane composition influences ABHD2 substrate preference.

Membrane trafficking may provide an additional level of regulation. Following agonist stimulation, ABHD2 and mPRα showed increased colocalization at the cell periphery at 30 min and within intracellular regions at 60 min, a pattern consistent with coordinated redistribution. Inhibition of endocytosis with Pitstop 2 attenuated OD-induced AMPK phosphorylation. These findings support a contribution of endocytic processes to mammalian mPRs signaling, extending our previous observations in *Xenopus* oocytes^11, 13, 16^, and suggest that membrane trafficking helps organize mPR-ABHD2 signaling across cellular compartments.

The metabolic experiments establish the functional importance of ABHD2 in mammalian mPRs signaling. ABHD2 deletion disrupted OD-induced AMPK activation, glycolysis and glucose uptake in HepG2 cells, as well as AMPK activation in C2C12 muscle cells. Re-expression of wild- type ABHD2 restored the glycolytic response, whereas the catalytic-site mutant did not. Together with the PLA₂ assays and our previous demonstration that OD-induced glycolysis depends on AMPK^8^, these findings support a model linking receptor-dependent ABHD2 catalytic activity to metabolic signaling. The intervening lipid mediators remain unknown. Lysophosphatidic acid and sphingosine-1-phosphate, previously implicated in progesterone signaling in oocytes^11^, are known to influence glucose uptake and hepatic glucose production^25, 26, 27^ and therefore represent candidate pathways for further investigation.

The cellular requirement for ABHD2 was accompanied by a significantly attenuated glucose- lowering response to OD in obese ABHD2 KO mice, establishing its contribution to the metabolic effects of mPRs agonism at the whole-body level. ABHD2 deletion also affected basal metabolism: knockout HepG2 cells exhibited elevated glycolysis and glucose uptake, while chow-fed knockout mice had lower fasting glucose levels. These findings suggest that ABHD2 contributes to basal glucose homeostasis as well as agonist responsiveness. Whether the basal phenotype reflects its physiological functions, compensatory adaptations to chronic deletion, or both remains unresolved.

In conclusion, our study identifies a conserved receptor-hydrolase partnership linking rapid steroid signaling to metabolic control. By directing ABHD2 localization and supporting its catalytic activity, mPRs couple hormone recognition to spatially organized enzymatic responses. These findings extend the mPR-ABHD2 partnership from amphibian oocyte maturation to mammalian metabolism and provide a broader framework for understanding how hormone-responsive protein complexes organize non-genomic steroid signaling.

## Methods

### Lead contact, materials and data availability

Requests for study-specific materials and supporting data should be directed to Nancy Nader. These are available upon request, with material transfers subject to applicable institutional agreements.

### mPRs agonist

The selective membrane progesterone receptor (mPR) agonist Org OD 02-0 (OD 02-0; Axon Medchem, Axon 2085) was used for all in vitro and in vivo studies.

### Cell lines

Human HepG2 hepatoma cells (ATCC HB-8065) and mouse C2C12 myoblasts (ATCC CRL- 1772) were obtained from ATCC and cultured according to the manufacturer’s recommendations. C2C12 cells were differentiated into myotubes in DMEM supplemented with 2% horse serum for 3–5 days. ABHD2-knockout HepG2 and C2C12 cells and mPRα-knockout HepG2 cells were generated using CRISPR/Cas9-mediated gene editing. Cells receiving a non-targeting guide served as controls. Genomic regions spanning the guide target sites were amplified by PCR and analyzed by Sanger sequencing. Loss of mPRα protein was assessed in two independent knockout clones by immunoblotting using an anti-mPRα antibody (Thermo Fisher Scientific, cat. PA5- 102102). Endogenous mPRβ expression was also assessed in naïve HepG2 cells using an anti- mPRβ antibody (Abcepta, cat. AP14849b). ABHD2 gene disruption in HepG2 and C2C12 cells was validated by Sanger sequencing; however, loss of endogenous ABHD2 protein could not be verified by immunoblotting because none of six commercial antibodies tested provided specific detection (Sigma-Aldrich, HPA005999; Proteintech, 140391-1-AP; Thermo Fisher Scientific, PA5-106585; antibodies-online, ABIN7160145; LSBio, LS-C101669; and Abcam, ab230417). Cells were routinely tested for mycoplasma contamination and confirmed to be mycoplasma-free.

### Expression constructs

Human ABHD2, human ABHD2 mutant (S207A/D345A/H376A) (ABHD2 mut), human mPRα, and human mPRβ expression constructs were generated by Gene Universal. For mammalian expression, coding sequences were cloned into pcDNA3 with fluorescent or epitope tags as indicated. Constructs included ABHD2-His, mPRα-GFP, mPRα-His, and ABHD2 fused to mCherry through an extended flexible linker at its C terminus, referred to as ABHD2-L-mCherry within the manuscript. Human Orai1 tagged with mCherry or GFP used as a plasma-membrane reference, are described previously ^28^. For cell-free expression, wild-type and catalytic-site mutant coding sequences were cloned into pALiCE02, which encodes an N-terminal melittin signal peptide for microsomal targeting and a C-terminal His tag.

### Transient transfection

HepG2 cells were transiently transfected using Lipofectamine 2000 Transfection Reagent (Invitrogen, cat. 11668019), according to the manufacturer’s instructions. Constructs were expressed individually or together in naive or knockout cells, as specified in the relevant experimental sections.

### High-resolution fluorescence microscopy experiments

HepG2 cells expressing the indicated ABHD2, mPRα, and Orai1 constructs were treated with vehicle or OD and examined by confocal or total internal reflection fluorescence (TIRF) microscopy. Imaging was performed in live or fixed cells, as specified in the figure legends. Human mCherry-tagged Orai1 served as a plasma-membrane reference where indicated. Nuclei were stained with Hoechst 33342 (Life Technologies, cat. H3570). For fixed-cell preparations, cells were incubated in 4% paraformaldehyde in PBS for 10 min at room temperature, washed with PBS, and permeabilized with 0.3% Triton X-100 in PBS for 10 min at room temperature. Following blocking with 3% Goat Serum and 1% bovine serum albumin in PBS for 1 hour at room temperature, cells were incubated with anti-His (LS-BIO, cat. LS-C60077) overnight at 4 °C. After washing, cells were incubated with Alexa Fluor 488-conjugated goat anti-rabbit IgG (abcam, cat. ab150081) for 1 h at room temperature, protected from light, and washed cells were then stained with Hoechst nuclear stain (Life technologies, cat. H3570) for 15 min at room temperature, protected from light, followed by washing with PBS before imaging.

*Confocal microscopy.* Confocal imaging was performed using a Zeiss LSM880 confocal microscope. To examine ABHD2 localization, ABHD2-deficient HepG2 cells expressing ABHD2- His were co-transfected with mCherry-tagged Orai1 as a plasma-membrane reference. Cells were treated with vehicle or OD, and ABHD2 was detected by anti-His (LS-BIO, cat. LS-C60077) immunostaining followed by an Alexa Fluor 488-conjugated secondary antibody (abcam, cat. ab150081). Images were acquired using matched settings across treatment groups within each experiment. Fluorescence line-scan analysis was performed in ImageJ. Lines were drawn from the extracellular region into the cell interior, crossing the plasma membrane, and fluorescence intensity profiles were extracted for the Orai1 and ABHD2 channels along the same line. Each profile was normalized to its own maximum intensity. The absolute distance between the Orai1 and ABHD2 fluorescence peaks was calculated and compared between vehicle- and OD-treated cells, with smaller distances indicating closer alignment of ABHD2 with the Orai1-defined plasma membrane. For colocalization experiments, naïve HepG2 cells co-expressing mPRα-His and ABHD2-L-mCherry were examined after vehicle treatment or 30 and 60 min of OD exposure. mPRα-His was detected by anti-His immunostaining, while ABHD2 was visualized through its mCherry fluorescence. Colocalization between mPRα-His and ABHD2-L-mCherry was quantified using Pearson’s correlation coefficient (PCC), calculated with the BIOP JACoP plugin in ImageJ. Regions of interest were selected separately at the cell periphery, guided by peripheral mPRα-His staining, and within the cytoplasm, excluding the nucleus and peripheral region. Each ROI was applied to both fluorescence channels. Consistent ROI selection criteria and analysis settings were used across treatment groups.

*Total internal reflection fluorescence microscopy (TIRF).* Live- and fixed-cell TIRF imaging was performed using a Zeiss microscope. ABHD2 was visualized using a fluorescent tag or anti-His immunostaining, with fluorescent Orai1 serving as a plasma-membrane reference. Time-lapse recordings were acquired to characterize basal ABHD2 dynamics. OD-induced ABHD2 accumulation was assessed in vehicle- and OD-treated cells, including ABHD2- and mPRα-KO cells. mPRα dynamics were examined in mPRα-KO cells re-expressing mPRα tagged with GFP. For fluorescence quantification, TIRF illumination was adjusted for both channels, and the Orai1 signal was used to identify the plasma-membrane imaging plane. Images of Orai1 and ABHD2 were acquired at this same focal plane. ABHD2 fluorescence intensity was quantified in ImageJ within the corresponding cell regions and compared between vehicle- and OD-treated cells.

### Co-immunoprecipitation assay

ABHD2-deficient HepG2 cells expressing His-tagged ABHD2 were treated with vehicle or OD for 30 min. Cells were lysed, and His-tagged ABHD2 and associated proteins were isolated using a His-tag magnetic pull-down kit (Anti-His MicroBeads, Miltenyi Biotec, Cat. No. 130-091-124), according to the manufacturer’s instructions. Eluted fractions were analyzed by immunoblotting for endogenous mPRα, endogenous mPRβ, and His-tagged ABHD2.

### Cell-free protein expression and microsomal preparations

Human ABHD2, human ABHD2 mutant (S207A/D345A/H376A) (ABHD2 mut), mPRα, and mPRβ were expressed individually or together using the ALiCE® Cell-Free Protein Expression Kit (Sigma-Aldrich), according to the manufacturer’s instructions. This tobacco BY-2 lysate-based system contains endogenous microsomal membranes that support protein translocation and membrane insertion. DNA was added to the cell-free lysates and incubated for 48 h at 25 °C with shaking at 700 rpm. The resulting microsomal preparations were used for progesterone-binding and PLA₂ activity assays without detergent extraction. Expression of wild-type and catalytic-site mutant proteins was verified by immunoblotting using an anti-His antibody (Cat. 631212, Takara).

### Progesterone binding assay

Human His-tagged ABHD2 and mPRα were expressed separately using the ALiCE cell-free protein expression system. Expression reactions were diluted 1:200 in binding buffer containing 30 mM HEPES and 100 mM NaCl (pH 7.5), and 20 µL was retained as input. The remaining samples were divided into 490-µL aliquots and incubated with progesterone-3-PEG11-biotin (Cayman Chemical, Cat. No. 9000645) at 10⁻⁵ M, or ethanol vehicle for 30 min. Pierce Streptavidin Magnetic Beads (Thermo Fisher Scientific, Cat. No. 88817) were blocked with 3% BSA and 0.2% Tween-20 for 30 min, washed with binding buffer, and incubated with the samples for 30 min. Beads were then washed sequentially with: (i) 1 M NaCl, 1% Ecosurf EH-9, 0.5% sodium deoxycholate, 0.1% SDS, and 50 mM Tris-HCl (pH 8.0) for 15 mins; (ii) 0.5 M NaCl, 30 mM HEPES, and 0.2% Triton X-100; and (iii) 20 mM Tris-HCl (pH 7.5). For displacement experiments, beads were incubated with free P4 or OD (10⁻⁴ M) for 1 h at room temperature following P4-biotin capture and washing. The competitor-containing supernatants were then removed, and proteins remaining on the beads were eluted in 2× Laemmli buffer and analyzed by Western blotting.

### PLA₂ activity in HepG2 cell lysates

Control, ABHD2-deficient and mPRα-deficient HepG2 cells were treated with vehicle or OD (10^-7^M) for the indicated durations. Cells were then lysed in RIPA Buffer, and 30µg of protein loaded per well in black 96-well plates with each condition loaded in three wells. PLA₂ activity was then measured using a fluorometric assay kit (Abcam, ab273278), according to the manufacturer’s instructions. Fluorescence was recorded over time, and background signal was subtracted before calculating activity. For each independent experiment, the maximal PLA₂ activity observed across the OD treatment time points was used for the summary analysis.

### PLA₂ activity in the reconstituted system

PLA₂ activity was assessed in microsomal preparations generated using the ALiCE cell-free expression system with a fluorometric PLA₂ assay kit (Abcam). Microsomes expressing wild-type ABHD2 (ABHD2 WT), the catalytic mutant ABHD2 (ABHD2 mut), mPRα, or mPRβ individually, or ABHD2 WT or ABHD2 mut co-expressed with either mPRα or mPRβ, were analyzed. Microsomal preparations were diluted 1:250 in assay buffer and were treated with progesterone (P4; 10⁻⁵ M), or ethanol as a vehicle control. Fluorescence was monitored kinetically for 60 min using a CLARIOstar microplate reader (BMG LABTECH). Expression of the relevant proteins was verified by immunoblotting.

### Effect of endocytosis inhibition on mPR-mediated signaling

#### Transferrin uptake assay

HepG2 cells were serum-starved in serum-free DMEM for 30 min at 37 °C, then pretreated with 20 µM Pitstop 2 (Abcam, cat. ab120687) or with vehicle (0.1% DMSO) for 40 min. The treatment medium was then removed and replaced with serum-free DMEM containing 50 µg/mL Alexa Fluor 488-conjugated human transferrin (Invitrogen, cat. T13342; stock prepared at 5 mg/mL in water). Following incubation for 15 min at 37 °C, cells were washed three times with ice-cold PBS and fixed with 4% paraformaldehyde for 10 min at room temperature. Nuclei were then stained with Hoechst. Fluorescence images were acquired using a Zeiss microscope.

#### Western blot analysis

HepG2 cells were pretreated with 20 µM Pitstop 2 for 40 min. Cells were then washed and treated with OD (10⁻⁷ M) or vehicle (ethanol) for the indicated times. Following treatment, cells were lysed and signaling responses were assessed by western blotting as described below.

### Signaling studies, protein extraction and immunoblotting

HEPG2 cells were maintained in growth medium consisting of DMEM supplemented with 20% heat-inactivated fetal bovine serum and 1% penicillin-streptomycin. C2C12 cells were cultured in growth medium (DMEM + 10% FBS + 1% Pen/Strep) for the first 24 hours, followed by differentiation in DMEM supplemented with 2% horse serum and 1% Pen/Strep for 3-5 days, until the formation of multinucleated myotubes was observed. After Overnight starvation, HepG2 and C2C12 myotubes were treated with ethanol (0.1%, as vehicle control) and OD 02-0 (10^-7^M) for 5, 10, 20 and 30 min. Cells were lysed in RIPA buffer and quantified as described above. Equal amounts of protein (30 μg) were separated by SDS–PAGE and transferred to PVDF membranes. Primary antibodies included phospho-AMPK (Cell Signaling Technology, cat. 2535S), total AMPK (Cell Signaling Technology, cat. 2532S), α-tubulin (Cell Signaling Technology, cat. 3873), β-actin (Cell Signaling Technology, cat. 3700), and GAPDH (abcam, cat. ab8245) were used as loading controls. Secondary antibodies included HRP-conjugated goat anti-rabbit IgG (Jackson ImmunoResearch, cat. 111-035-144), HRP-conjugated goat anti-mouse IgG (Jackson ImmunoResearch, cat.115-035-003), infrared-conjugated goat anti-rabbit IgG (LI-COR, cat. 925- 32211) and goat anti-mouse IgG (LI-COR, cats. 925-68070 and 926-32210). Immunoblot signals were detected using ECL Prime Western Blotting Detection Reagent (GE Healthcare Amersham, cat. 45-002-401) or a LI-COR Odyssey imaging system. Phosphorylated and total AMPK signals were normalized to the loading control and expressed as a phospho-to-total ratio, followed by normalization to the comparator specified in the figure legend.

### Extracellular flux analysis - Glycolysis stress test

Glycolytic activity was assessed by measuring extracellular acidification rate (ECAR) using a Seahorse XF Glycolysis Stress Test Kit (Agilent, cat. 103020-100). HepG2 cells were seeded at 30,000 cells per well in Seahorse XF96 plates, with each plate containing a single genotype or rescue condition. Half of the wells were treated with vehicle (0.1% ethanol) and half with OD (10⁻⁶ M) for 2 h before analysis. ECAR was measured at baseline and following sequential injections of glucose, oligomycin and 2-deoxyglucose. Data were analyzed using Seahorse Wave software (Agilent). Glycolysis was calculated as the maximum ECAR after glucose addition and before oligomycin injection minus the last ECAR measurement before glucose addition. Each full plate represented one independent experiment, with wells serving as technical replicates. Technical replicate values were averaged for each condition, and glycolysis was expressed relative to the matched vehicle-treated control within each experiment.

### Intracellular glucose measurements by FRET imaging

Intracellular glucose responses were monitored using the genetically encoded CFP/YFP sensor FLII¹²Pglu-700μδ6. The sensor coding sequence from pcDNA3.1-FLII12Pglu-700uDelta6 (Addgene, plasmid 17866) was inserted into the BglII/HindIII sites of pShuttle-CMV (Addgene, plasmid 16403) for adenoviral expression. HepG2 cells were infected at a multiplicity of infection of 100. Following vehicle or OD treatment for 30 min, cells were challenged with 1 mM glucose. Live-cell imaging was performed using a Zeiss LSM880 confocal microscope with 458-nm excitation and CFP and YFP detection at the reported wavelengths of 482 and 570 nm, respectively. YFP/CFP ratios were calculated using ImageJ and normalized to the baseline before glucose addition. Glucose uptake (ΔFRET) was calculated from the raw ratio data by subtracting the ratio immediately before glucose addition from that measured after glucose addition.

### Mouse models of diet-induced obesity and treatment

Age-matched male C57BL/6N mice were obtained from Jackson Laboratory, and ABHD2 KO mice on a C57BL/6N background were generously provided by Dr. Polina Lishko (Washington University)^29^. All animal experiments were approved by the Weill Cornell Medicine-Qatar Institutional Animal Care and Use Committee. Mice were housed individually in ventilated cages in a specific pathogen-free facility, with standard bedding and environmental enrichment, under a 12-h light–dark cycle at 22 ± 2°C and 50–60% relative humidity. Food and water were available ad libitum except during fasting, and animal health was monitored regularly by trained staff. At approximately 20 weeks of age, mice were maintained on chow or fed a high-fat diet providing 60% of calories from fat (Research Diets) for up to 17 weeks. Body weight was recorded weekly, and blood glucose was measured after a 6-h fast using an AlphaTrak 3 glucometer. Following confirmation of HFD-induced hyperglycemia, mice were randomized to treatment groups balanced for mean pretreatment fasting glucose. Mice received OD 02-0 (4 mg/kg/day) or vehicle containing 10% hydroxypropyl-β-cyclodextrin (HP-β-CD) and 10% ethanol by oral gavage for four weeks. OD was initially dissolved in ethanol at 10 mg/mL, and the amount added to each dose was adjusted according to body weight while maintaining a final gavage volume of 200 µL and the same vehicle composition. Fasting blood glucose was reassessed after treatment.

## Statistical analysis

Data are presented as mean ± SEM. Statistical analyses were performed using GraphPad Prism 10 with the tests specified in the figure legends. Statistical significance is denoted as follows: ns, not significant; *p ≤ 0.05; **p ≤ 0.01; ***p ≤ 0.001; and ****p ≤ 0.0001.

## Acknowledgments

We are grateful for the Vivarium and microscopy core at WCMQ for their support in multiple experiments. We also thank Dr. Polina Lishko for generously providing the ABHD2 knockout mice. The authors used ChatGPT (OpenAI, GPT-5.2) for grammar and language refinement. This work was supported by the Biomedical Research Program at Weill Cornell Medical College in Qatar (BMRP) awarded to Nancy Nader, a program funded by Qatar Foundation. The statements made herein are solely the responsibility of the authors.

## Contribution statement

N. Nader conceived and designed the study, secured funding, supervised the project, led data analysis and interpretation, and wrote the manuscript. S. Sherif, L. Assaf, R. El Ghali, Mariam A., and M. Shammout performed experiments and acquired data. K. Machaca contributed to scientific discussions. All authors revised the manuscript and approved the final version. N. Nader is the guarantor of this work and, as such, has full access to all the data in the study and takes responsibility for the integrity of the data and the accuracy of the data analysis.

---

**Fig. S1.** Generation of ABHD2-KO HepG2 cells and constructs used to examine ABHD2 localization. (A) Sequence chromatograms spanning the ABHD2 CRISPR/Cas9 target site in non-targeting control HepG2 cells and clones T2-32 and T2-44. The guide RNA sequence is AATTCCAGGGCAGATGACCA. The target region is highlighted in black, and arrows indicate insertion/deletion sites. **(B)** Predicted ABHD2 amino acid sequences in clones T2-44. Predicted stop codons are highlighted in yellow. **(C)** Schematic of the ABHD2-(GGGS)₅-mCherry construct, with a flexible linker between the ABHD2 C terminus and mCherry (ABHD2-L-mCherry). **(D)** Representative confocal image of ABHD2-KO HepG2 cells expressing ABHD2-His detected by anti-His immunostaining showing intracellular puncta and track-like structures. The boxed region is enlarged fourfold at right. Scale bar, 20 μm.

**Fig. S2.** Validation and functional characterization of mPRα-KO HepG2 cells. **(A)** Sequence chromatograms spanning the PAQR7 CRISPR/Cas9 target site in non-targeting control HepG2 cells and clones T2-15 and T2-16. The guide RNA sequence is AAGCGCCAGGTCTGATGCAG. The target region is highlighted in black, and arrows indicate insertion/deletion sites. **(B)** Predicted mPRα amino acid sequences in clones T2-15, with predicted stop codons highlighted in yellow. **(C)** Representative immunoblots of mPRα in parental (naïve), non-targeting control (Neg. Ctrl), and PAQR7-targeted HepG2 clones T2-15 and T2-16. Tubulin served as the loading control. **(D)** Representative immunoblots (left) and quantification (right) of AMPK phosphorylation in mPRα-KO clone T2-15 treated with EtOH (0.1%) or OD (10⁻⁷ M) for 5 to 30 min. Tubulin served as the loading control. Phospho-AMPK/Tubulin (R1) and total-AMPK/Tubulin (R2) ratios were used to calculate R1/R2, followed by normalization to EtOH. The OD treatment time point yielding maximal activity in each independent experiment was plotted. Data are presented as mean ± SEM; n=5 independent experiments. Values between ethanol and OD treatment were compared using unpaired *t* test. **(E)** Schematic of the FLII¹²Pglu-700μδ6 glucose sensor and representative CFP, YFP, and merged fluorescence images of sensor-expressing HepG2 cells. **(F)** Normalized FRET ratio following addition of 1 mM glucose to mPRα-KO clone T2-15 pretreated with EtOH (0.1%) or OD (10^-6^M) for 30 min (left). Right, ΔFRET quantification before normalization. Data are presented as mean ± SEM; n = 27 to 29 cells per condition from 3 independent experiments. Values between ethanol and OD treatment were compared using unpaired *t* test. **(G)** Representative fixed-cell TIRF images of mPRα-KO HepG2 cells expressing ABHD2-His (green) and Orai1-mCherry (red), following EtOH or OD treatment for 30 min. Quantification is included in Fig. 1G. Statistical significance: *p < 0.05; ns, not significant.

**Fig. S3.** progesterone association with human ABHD2 and mPRα **(A)** Anti-His immunoblot of ALiCE preparations expressing human ABHD2 or mPRα. Unprogrammed lysate is shown as a control. **(B)** Progesterone-biotin pull-downs of human ABHD2 (left) and mPRα (right) expressed separately in the ALiCE system. Preparations were incubated with progesterone-3-PEG11-biotin (10⁻⁵ M) or EtOH (0.1%) and recovered using streptavidin beads. For competitive elution, captured proteins were incubated with free P4 or OD (10⁻⁴ M). Bound/eluted proteins were detected by anti- His immunoblotting. **(C)** Immunoblot showing endogenous mPRβ expression in HepG2 cells. Actin served as the loading control.

**Fig. S4.** mPRβ supports catalytic-site-dependent PLA₂ activity of human ABHD2. **(A)** anti-His immunoblot confirming expression of human mPRα, mPRβ, WT ABHD2, and catalytic-site mutant ABHD2 (ABHD2 mut), individually or in the indicated combinations, in ALiCE preparations. **(B)** Time courses of background-corrected PLA₂ substrate hydrolysis in microsomal preparations expressing mPRβ, WT ABHD2, or catalytic-site mutant ABHD2, individually or together as indicated, in the presence or absence of P4 (10⁻⁵ M). EtOH served as vehicle. **(C)** PLA₂ activity at 30 min in microsomes preparations co-expressing mPRβ with WT or catalytic- site mutant ABHD2, in the presence or absence of P4. Data are presented as mean ± SEM, n=6. Values between the different conditions were compared using one-way ANOVA. Statistical significance: ****p < 0.0001; ns, not significant.

**Fig. S5.** Endocytosis perturbation alters OD-induced signaling **(A)** Live-cell TIRF images of mPRα KO HepG2 cells expressing mPRα-GFP (green) together with Orai-mCherry (red). Scale bar, 5 μm. **(B)** Confocal images of Alexa Fluor 488-transferrin uptake (green) in HepG2 cells pretreated with vehicle (0.1% DMSO) or Pitstop 2 (20 μM) for 40 min, followed by incubation with transferrin (50 μg/mL) for 15 min. Nuclei were stained with Hoechst (blue). **(C, D)** Representative immunoblots (C) and quantification (D) of phospho- and total AMPK following EtOH or OD treatment (10⁻⁷ M, 5 to 30 min), after pretreatment with Pitstop 2 (20 μM, 40 min), followed by washing. GAPDH served as loading controls. Phospho-AMPK/GAPDH (R1) and total-AMPK/GAPDH (R2) ratios were used to calculate R1/R2, followed by normalization to EtOH. The OD treatment time point yielding maximal activity in each independent experiment was plotted. Data are presented as mean ± SEM; n=3 independent experiments. Values between ethanol and OD treatment were compared using unpaired *t* test. Statistical significance: ***p < 0.001; ns, not significant.

**Fig. S6.** Basal metabolic effects of ABHD2 deletion in HepG2 cells and loss of OD-induced AMPK activation in C2C12 cells. **(A)** Basal AMPK phosphorylation in negative control (Neg. Ctrl) compared to ABHD2-KO HepG2 cells, expressed as phospho-AMPK/Tubulin (R1) divided by total-AMPK/Tubulin (R2). Data are presented as mean ± SEM; n = 3 to 5 independent experiments. Values between the 2 conditions were compared using unpaired *t* test. **(B)** Quantification of glycolysis in vehicle-treated (ethanol 0.1%) negative-control (Neg. Ctrl) and ABHD2-KO HepG2 cells. Data are mean ± SEM; n = 3. Groups were compared using an unpaired *t* test. **(C)** ΔFRET in negative control (Neg. Ctrl) AND ABHD2-KO HepG2 cells in vehicle condition (ethanol 0.1%). Raw ΔFRET were calculated as the increase in the YFP/CFP ratio without normalization after glucose addition relative to the ratio immediately before addition. Data are presented as mean ± SEM from 3 independent experiments. Values were compared using unpaired *t* test. **(D)** Sequence chromatograms spanning the ABHD2 CRISPR/Cas9 target site in non-targeting control C2C12 cells and clones T2-8 and T2-16. The guide RNA sequence is CCGCAAGTTCATCACCATGT. The target region is highlighted in black, and arrows indicate insertion/deletion sites. **(E)** Predicted ABHD2 amino acid sequences in the indicated C2C12 KO clones, with predicted stop codons highlighted in yellow. **(F, G)** Representative immunoblots (left) and quantification (right) of AMPK phosphorylation in differentiated non-targeting control (Neg. ctrl) (F) and ABHD2-KO (G) C2C12 cells treated with EtOH (0.1%) or OD (10⁻⁷ M) for 5 to 30 min. GAPDH served as the loading control. Phospho- AMPK/GAPDH (R1) and total-AMPK/GAPDH (R2) ratios were used to calculate R1/R2, followed by normalization to EtOH. Data are presented as mean ± SEM; n = 3 independent experiments. Values between Ethanol and OD were compared using unpaired *t* test. Statistical significance: **p < 0.01; ***p < 0.001; ns, not significant.

**Fig. S7.** ABHD2 deletion alters the glucose-lowering effect of OD in obese mice. **(A, D)** Fasting blood glucose after a 6-h fast (A) and body weight (D) in chow-fed male C57BL/6N (n=35) and ABHD2-KO mice (n=66). Data are presented as mean ± SEM. Fasting glucose and body weight values between C57BL/6N and ABHD2-KO mice were compared using Welch’s *t* test. **(B, C, E, F)** Fasting blood glucose (B, C) and body weight (E, F) before and after high-fat diet (HFD) feeding in C57BL/6N mice (B, E; n = 10 for fasting glucose and 6 for body weight) and ABHD2-KO mice (C, F; n = 15 for fasting glucose and 14 for body weight). Data are presented as mean ± SEM. Before- and after-HFD values were compared using paired *t* tests. **(G, H)** Fasting blood glucose after a 6-h fast right before and after 4 weeks of OD treatment (4 mg/kg/day by oral gavage) in HFD-fed C57BL/6N (n=6) (G) and ABHD2-KO (n=7) (H) mice. Lines connect measurements before and after treatment from the same mouse. Data are presented as mean ± SEM; Fasting glucose before and after treatment were compared using the paired *t* test. **(I)** Individual treatment-associated changes in fasting glucose, calculated as after-treatment minus before-treatment glucose for each mouse. Negative values indicate a decrease. Each point represents one mouse. Data are presented as mean ± SEM; C57BL/6N (n=6); ABHD2 KO (n=7). The dip in glucose values between C57BL/6N and ABHD2-KO mice were compared using Welch’s *t* test. **(J)** Percentage Body-weight gain over time in male ABHD2-KO mice maintained on chow or high- fat diet (HFD), followed by treatment with vehicle (HFD-V) or OD (HFD-OD; 4 mg/kg/day by oral gavage) for 4 weeks. The percentage gain is calculated based on the starting body weight. Arrows indicate HFD initiation and treatment initiation. Data are presented as mean ± SEM; n = 7 mice chow, n=6 mice HFD-V, n=7 mice HFD-OD. Statistical significance: *p < 0.05; **p < 0.01; ***p < 0.001; ****p < 0.0001; ns, not significant.

## Supplementary Video 1- ABHD2 dynamics near the plasma membrane

Representative live-cell TIRF recording of ABHD2-KO HepG2 cells co-expressing ABHD2-L- mCherry and the plasma membrane marker Orai1-GFP under basal conditions.

**(A)** The ABHD2-L-mCherry channel is shown alone to facilitate visualization of dynamic membrane-proximal structures.

**(B)** Merged ABHD2-L-mCherry (red) and Orai1-GFP (green) channels from the same recording. Images were acquired using a 63×/1.46 NA oil-immersion objective at approximately 1.85-s intervals. Movies were exported in AVI format at 15 frames per second for accelerated playback.

## Supplementary Video 2 - ABHD2 dynamics below the plasma membrane

Representative live-cell TIRF recording showing dynamic ABHD2-L-mCherry-positive structures (red) below the plasma membrane under basal conditions in ABHD2-KO HepG2 cells expressing ABHD2-L-mCherry. Images were acquired using a 63×/1.46 NA oil-immersion objective at approximately 1-s intervals. The time series was exported as an AVI movie at 15 frames per second for accelerated playback.

## Supplementary Video 3 - mPRα dynamics near the plasma membrane

Representative live-cell TIRF recording showing dynamic mPRα-GFP-positive structures (green) near the plasma membrane under basal conditions in mPRα-KO HepG2 cells re-expressing mPRα- GFP. Images were acquired using a 63×/1.46 NA oil-immersion objective at approximately 1.72- s intervals. The time series was exported as an AVI movie at 15 frames per second for accelerated playback.

