## Supplemental figures for "Membrane progesterone receptors spatially organize and partner with ABHD2 to mediate metabolic signaling"

#### Supplemental Figure 1

**A**

Guide RNA - AATTCCAGGGCAGATGACCA

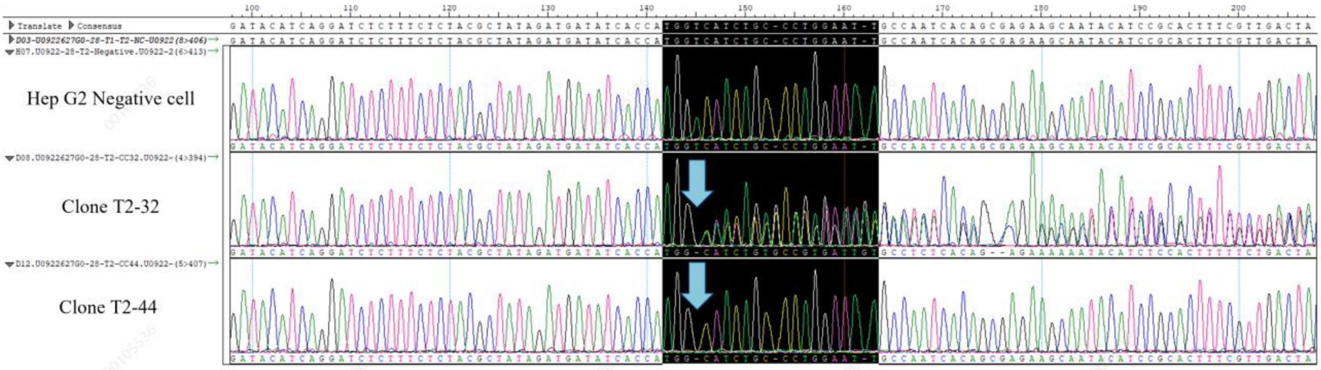


**B**

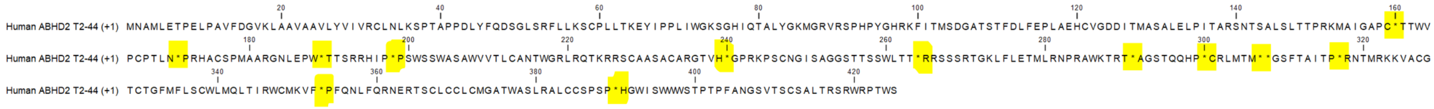


**C**

ABHD2–(GGGS)$_5$–mCherry

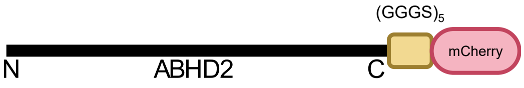


**D**

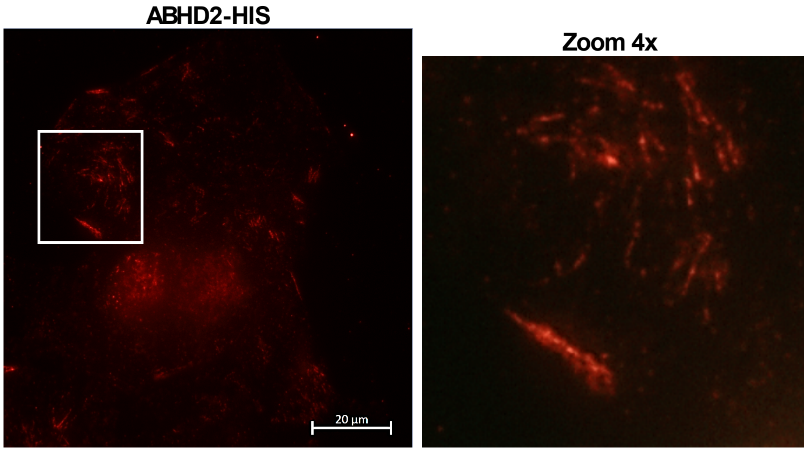

**Supplemental Figure 2**

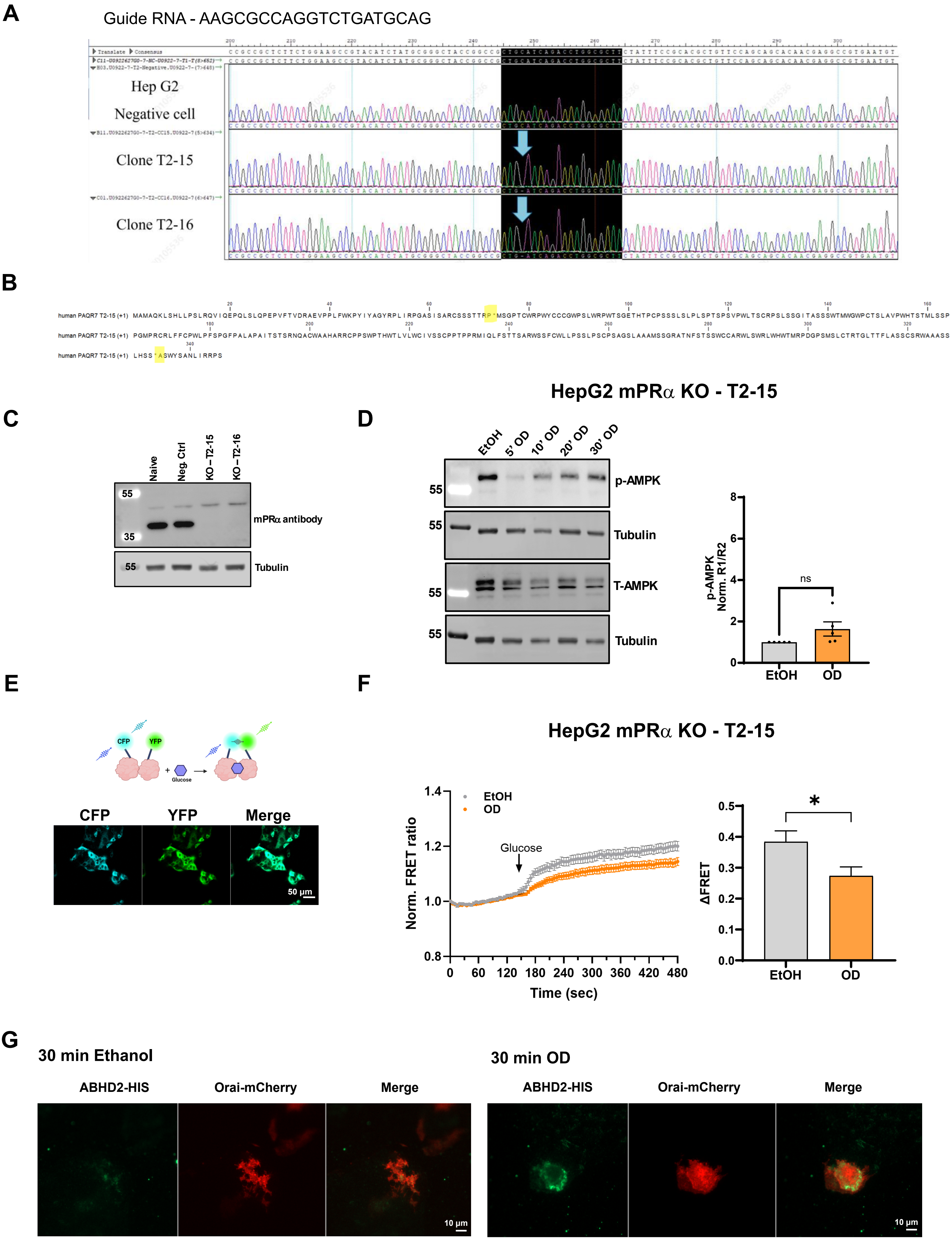

#### Supplemental Figure 3

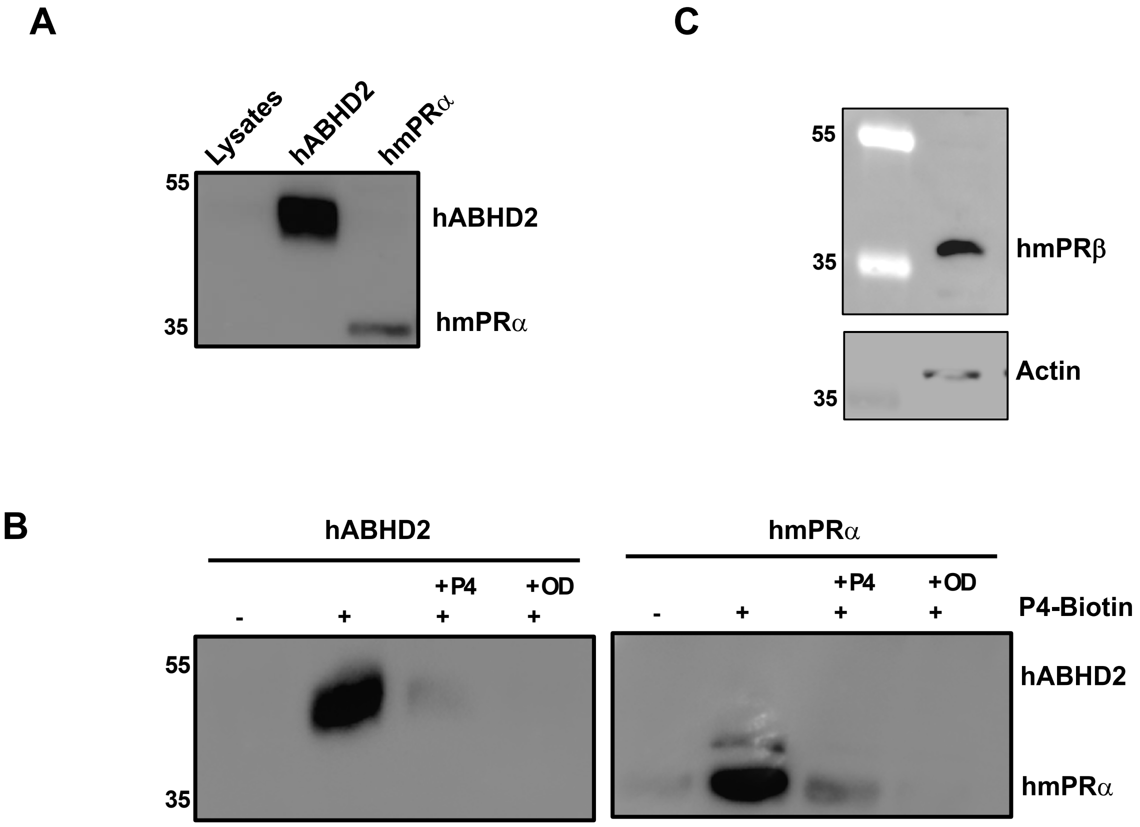

**Supplemental Figure 4**

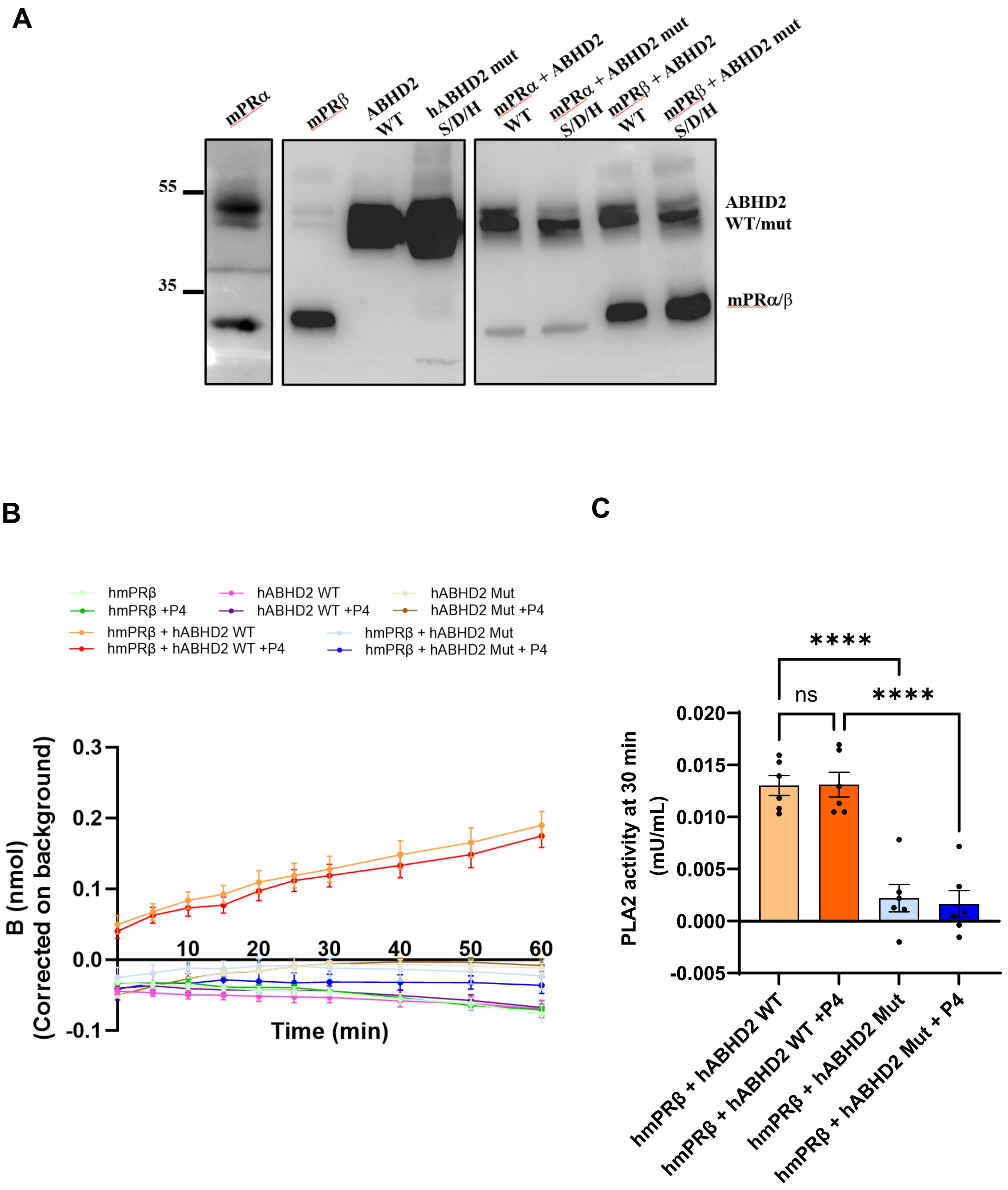

### Supplemental Figure 5

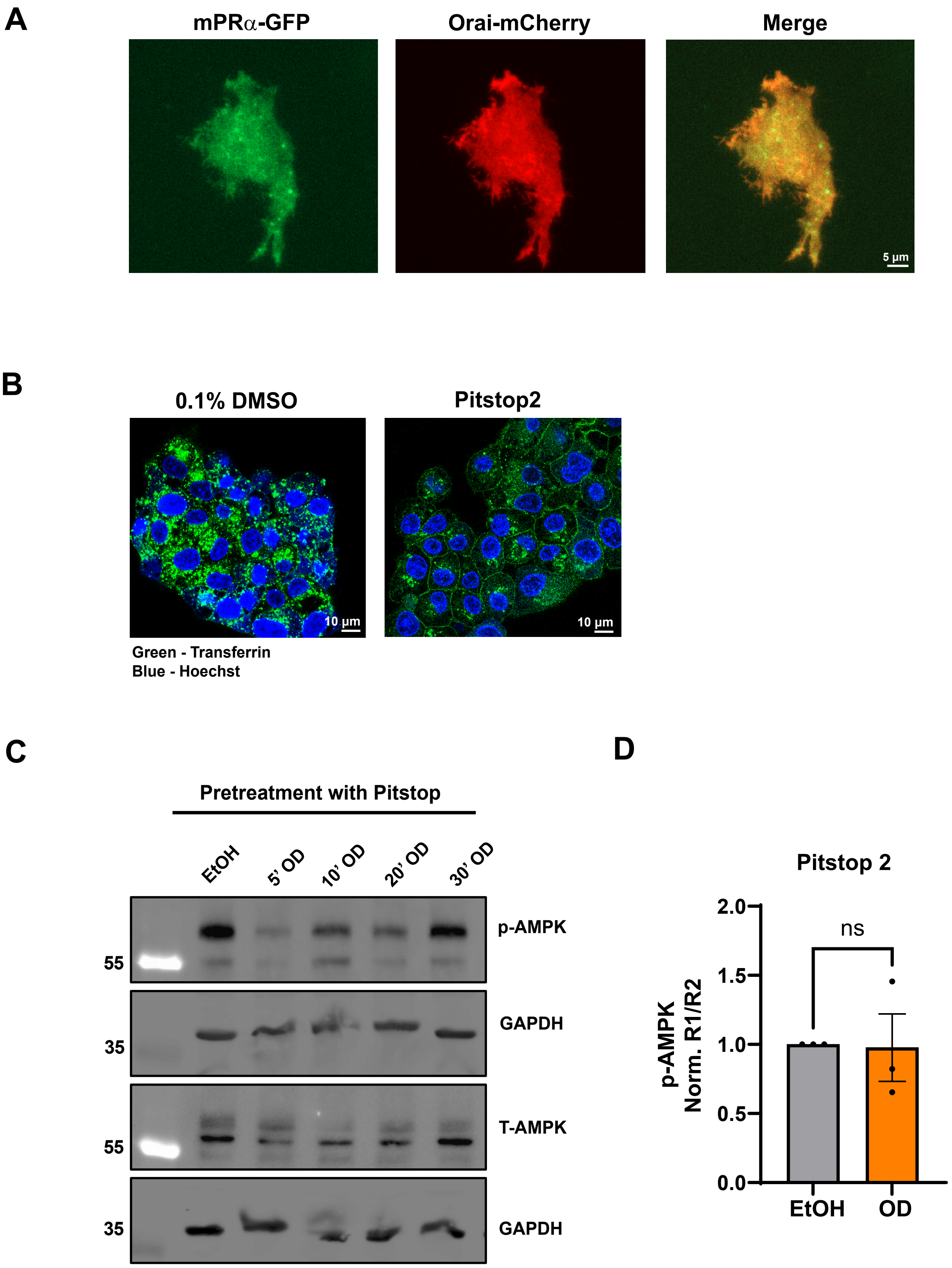

#### Supplemental Figure 6

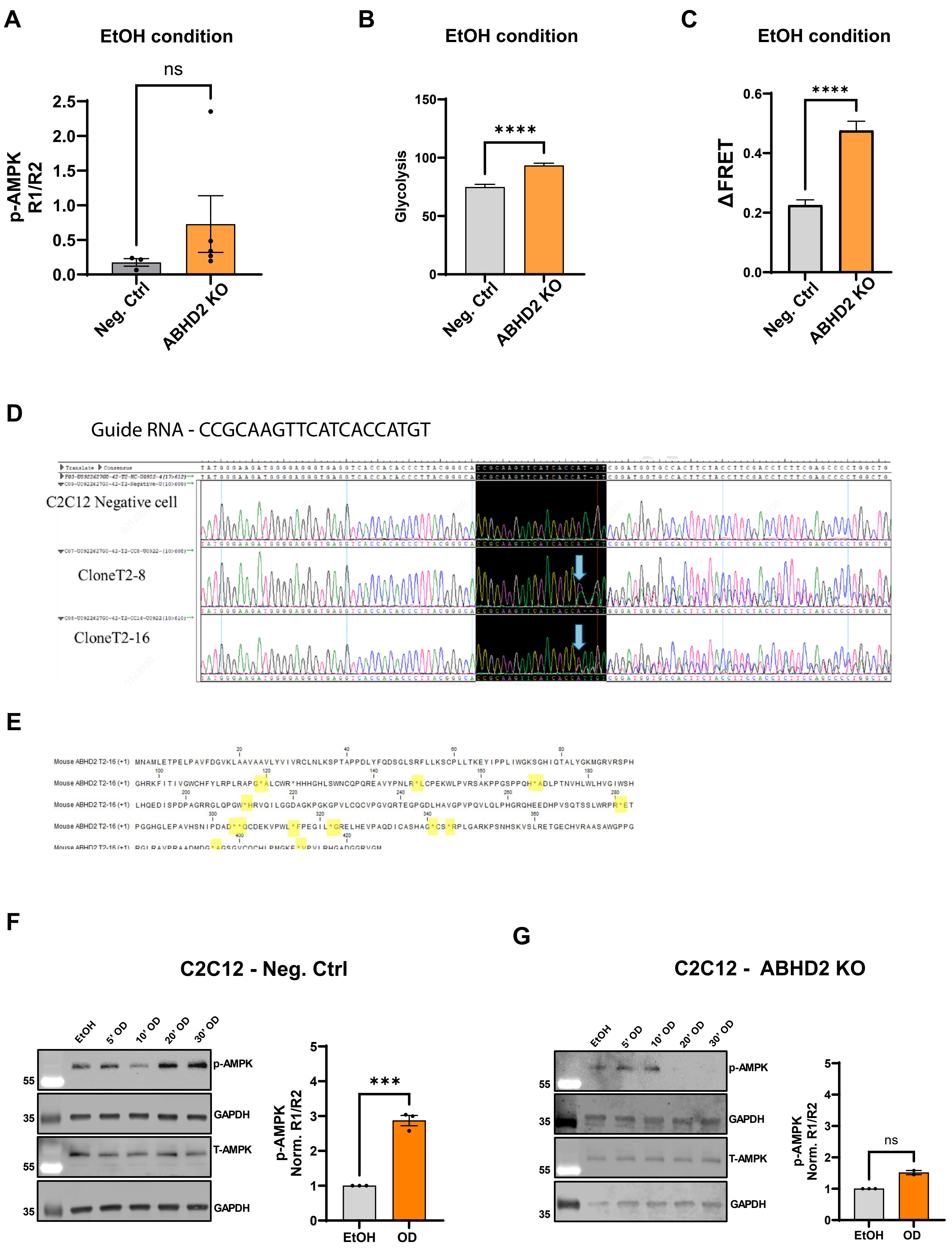

#### Supplemental Figure 7

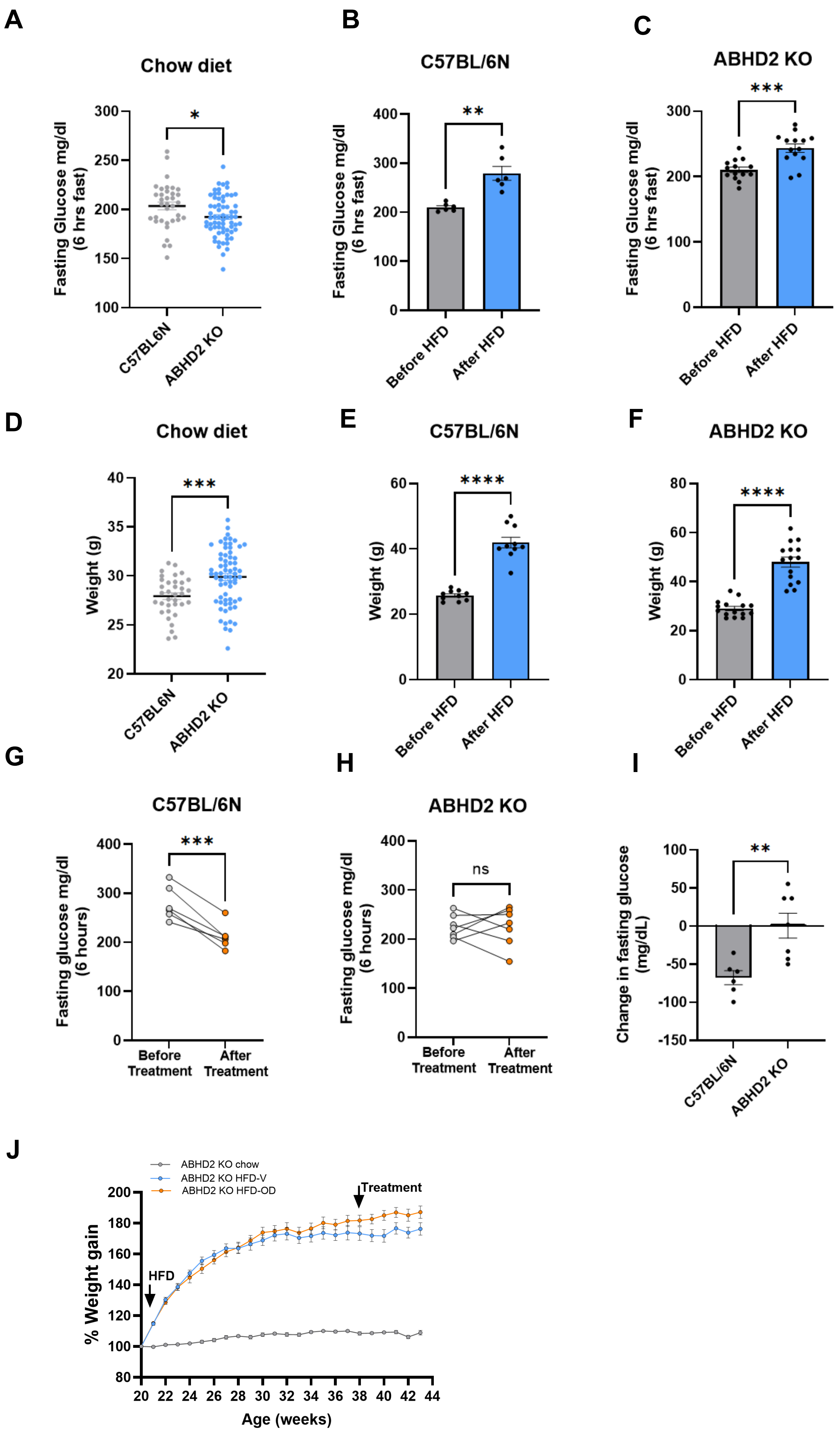
